# Visual mental imagery engages the left fusiform gyrus, but not the early visual cortex: a meta-analysis of neuroimaging evidence

**DOI:** 10.1101/2020.02.06.937151

**Authors:** Alfredo Spagna, Dounia Hajhajate, Jianghao Liu, Paolo Bartolomeo

## Abstract

The dominant neural model of visual mental imagery (VMI) stipulates that memories from the medial temporal lobe acquire sensory features in early visual areas. However, neurological patients with damage restricted to the occipital cortex typically show perfectly vivid VMI, while more anterior damages extending into the temporal lobe, especially in the left hemisphere, often cause VMI impairments. Here we present two major results reconciling neuroimaging findings in neurotypical subjects with the performance of brain-damaged patients: (1) a large-scale metaanalysis of 46 fMRI studies, of which 27 investigated specifically visual mental imagery, revealed that VMI engages fronto-parietal networks and a well-delimited region in the left fusiform gyrus. (2) A Bayesian analysis showing no evidence for imagery-related activity in early visual cortices. We propose a revised neural model of VMI that draws inspiration from recent cytoarchitectonic and lesion studies, whereby fronto-parietal networks initiate, modulate, and maintain activity in a core temporal network centered on the fusiform imagery node, a high-level visual region in the left fusiform gyrus.

## Introduction

Close your eyes and think of Leonardo da Vinci’s *Monna Lisa*. Is she looking at you or not? Is her hair curled or straight? Visual Mental Imagery (VMI) is the set of abilities whereby we can “see” things that are elsewhere (or nowhere: now imagine *Monna Lisa* frowning at you). These capacities are important for predicting the outcome of everyday tasks (Moulton and Kosslyn, 2009), for example to decide whether our car fits in a narrow parking spot. The subjective vividness of visual mental images varies substantially across individuals (Galton, 1880; Cui et al., 2007; Pearson et al., 2015; Dijkstra et al., 2017a), with some individuals experiencing mental images as “quasi-visual” in nature, while others having less vivid images, down to the total absence of VMI experience in otherwise normal individuals, a condition dubbed as “aphantasia” (Zeman et al., 2015; de Vito and Bartolomeo, 2016; Fulford et al., 2018; Jacobs et al., 2018). The neural bases of this remarkable set of cognitive functions are the object of intense research efforts (Ishai et al., 2000; Pearson et al., 2013; Dentico et al., 2014; Pearson et al., 2015; Dijkstra et al., 2017b; Dijkstra et al., 2018; Winlove et al., 2018). Identifying the brain circuits supporting VMI and motor imagery is also essential for clinical reasons, because detecting their activity in neuroimaging can reveal consciousness in non-communicating patients in apparent vegetative state (Owen et al., 2006); in addition, uncontrolled VMI activity could contribute to the vivid recollections of traumatic memories resulting in post-traumatic stress disorder (Mary et al., 2020).

The dominant model of VMI (Pearson et al., 2015; Dijkstra et al., 2019) stipulates the existence of common neural substrates underlying VMI and visual perception, spanning across the ventral cortical visual stream, with a crucial implication of early visual areas, providing the sensory and spatial representational content of VMI (Kosslyn et al., 2006; Pearson, 2019, 2020). Neuroimaging studies in healthy volunteers supported a strong version of the model, by demonstrating the engagement of early, occipital visual areas in VMI (Kosslyn et al., 2001;

Ibáñez-Marcelo et al., 2019). Further, TMS interference on V1 was shown to impact VMI (Kosslyn et al., 1999). The model also provided a principled account of inter-individual differences in VMI, because the level of activation in low-level visual areas correlates with the subjective experience of VMI “vividness” (Lee et al., 2012; Dijkstra et al., 2017a; but see Fulford et al., 2018). However, this neuroimaging evidence is correlational, and does not directly speak to the causal role of these structures in VMI. In fact, causal evidence from neurological patients is sharply discordant with an implication of early visual areas in VMI (Bartolomeo, 2002, 2008; Bartolomeo et al., 2013; Bartolomeo et al., 2020). Whereas the model would predict a systematic co-occurrence of perceptual and imaginal deficits after brain damage (Farah et al., 1988), patients with brain damage restricted to the occipital cortex often have spared VMI abilities (Behrmann et al., 1992; Chatterjee and Southwood, 1995; Policardi et al., 1996; Aglioti et al., 1999), with preserved subjective VMI vividness despite damaged early visual cortex (Goldenberg et al., 1995), even in the case of bilateral cortical blindness (Chatterjee and Southwood, 1995; Zago et al., 2010; de Gelder et al., 2015). Instead, deficits of VMI for object form, object color, faces or orthographic material typically arise as a consequence of more anterior damage (Bisiach and Luzzatti, 1978; Bisiach et al., 1979; Basso et al., 1980; Farah et al., 1988; Riddoch, 1990; Beschin et al., 1997; Manning, 2000; Sirigu and Duhamel, 2001), often extensively including the temporal lobes, especially in the left hemisphere (Bartolomeo, 2002, 2008; Moro et al., 2008). Such a strong evidence of dissociation is at odds with models proposing a crucial implication of early visual areas in VMI and suggests an engagement in VMI of higher-order associative areas, especially in the left temporal lobe, rather than early visual cortex. Also, growing evidence shows that VMI and perception build on distinct brain connectivity patterns (Dijkstra et al., 2017b), characterized by a reversed cortical information flow (Dentico et al., 2014), and supported by anatomical connectivity between striate, extrastriate, and parietal areas (Whittingstall et al., 2014). In addition, studies on VMI mainly focused on the ventral cortical visual stream, but evidence also indicates VMI-related increases in BOLD response in fronto-parietal networks (Mechelli et al., 2004; Yomogida et al., 2004; Mazard et al., 2005). Thus, what VMI shares with visual perception may not be the passive, low-level perceptual aspect, but the active, exploratory aspects of vision (Thomas, 1999; Bartolomeo, 2002; Bartolomeo et al., 2013), such as those sustained by visual attention (Bartolomeo and Seidel Malkinson, 2019). Yet, the precise identity of these mechanisms, and of the underlying brain networks, remains unknown.

To address these issues, we conducted a meta-analysis of functional magnetic resonance imaging studies that examined the neural correlates associated with visual and motor mental imagery. Our specific aims were to assess the role of low- and high-level visual cortex, as well as the role of the fronto-parietal networks in mental imagery, by identifying the brain regions with higher activation for mental imagery in healthy volunteers. Specifically, we asked whether or not VMI recruits early visual areas, defined as retinotopically organized visual areas (i.e., visual field maps according to e.g., Wandell and Winawer, 2011; Wang et al., 2015). To evaluate the relationships between modality-specific and more general processes in mental imagery, we also assessed the neuroimaging of *motor* mental imagery (MMI). The studies included the coordinates of brain regions with a significant activation increase in the contrasts of: VMI *versus* Control; VMI *versus* Perception; and MMI *versus* Control. We predicted the involvement in VMI of fronto-parietal and cingulo-opercular network areas, together with high-level areas in the visual cortical ventral stream. Based on evidence from clinical neurology, we did not expect to find an activation of the early visual areas associated with visual mental imagery. If confirmed, this prediction would challenge the current dominant model, postulating a crucial implication of early visual cortex in VMI, and delineate a new, alternative model, based on a critical role in our VMI experience of high-level visual region and of fronto-parietal networks important for attention and visual working memory.

## Method

### Literature search

First, we searched for functional magnetic resonance imaging (fMRI) studies in the domain of mental imagery using the online database NeuroSynth (http://NeuroSynth.org-RRID:SCR_006798) (Yarkoni et al., 2011). At the time of access (October 15, 2018), the database contained 3,884 activation peaks from 84 studies. To achieve an exhaustive search of studies, we further expanded our literature search to other databases (i.e., PubMed, Google Scholar), using the following search terms: Visual mental imagery, Mental imagery, Vividness, Mental rotation, and carefully screened the reference list of previously conducted reviews and meta-analyses. This final step led to the discovery of 39 additional studies.

### Inclusion and Exclusion criteria

We first applied the following inclusion criteria: (1) studies published in peer-review journals in English; (2) studies with normal healthy adults (age ≥ 18 years); (3) studies with table(s) reporting brain activation foci from whole-brain analysis; (4) studies with the coordinates of the activation reported either in the Montreal Neurological Institute (MNI) or in the Talairach space. We then applied the following exclusion criteria: (1) single-subject studies (excluded n = 2); (2) studies not using fMRI (e.g., voxel based morphometry studies or DTI studies; n = 8); (3) non-first hand empirical studies such as review articles (excluded n = 7); (4) studies of special populations of adults (e.g., with a specific personality trait or neurological disorders), whose brain functions may deviate from neurotypical controls’ (excluded n = 31), unless they reported also the results from the GLM or contrasts of interest; (5) studies not reporting coordinates for contrasts of interest in this study (n = 2). (6) studies that could not be clearly classified into one of our three subdomains, or reporting contrasts not of interest for this study (n = 19); (7) studies using analytic approaches not of interest for this study (e.g., ROI analyses, multivariate pattern analysis, functional or effective connectivity analysis) (n = 15), unless they reported also the results from the GLM or contrasts of interest. The resulting final number of studies included in the meta-analysis was 41.

### Data extraction

Within the included studies, we identified the tables that reported positive activation from defined contrasts of (1) VMI greater than control or rest conditions; (2) VMI greater than visual perception condition; (3) MMI greater than control or rest condition. “Control” conditions included experimental conditions unrelated to the specific imagery task. “Visual Perception” included all conditions in which participants viewed visual stimuli related to the items employed in the VMI conditions. In the present work, we will use the term “experiment” as the result of one statistical map, with a single study potentially reporting results from more than one map. In fact, five of the 41 studies reported multiple experiments or multiple contrasts of interest for this study (e.g., reporting coordinates for both VMI *versus* Control and VMI *versus* Perception or VMI *versus* Control and MMI *versus* Control), which brought the total number of included experiments to 46. Specifically, we found (1) 27 experiments (376 foci from a total of 380 subjects) reporting contrasts regarding the difference in activation between VMI and rest or control condition; (2) 4 experiments (62 foci from a total of 52 subjects) reporting contrasts regarding the difference in activation between visual mental imagery and perception, and (3) 15 experiments (340 foci from a total of 239 subjects) reporting contrasts regarding differences in activation between motor mental imagery and rest. **Figure 1** illustrates the selection process based on inclusion and exclusion criteria.

**Figure 1.**
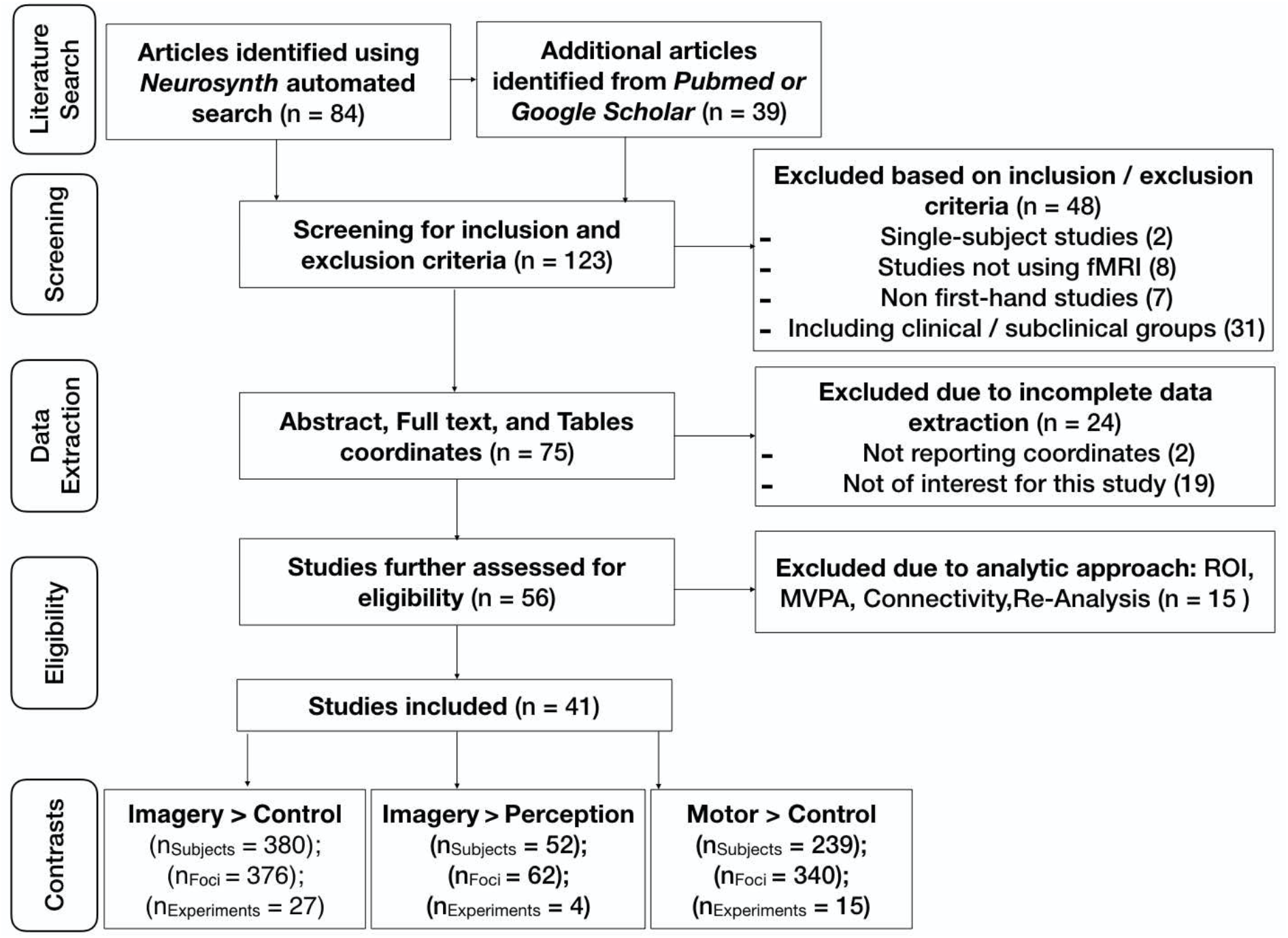
Flow chart of the selection process conducted in the present meta-analysis showing the following steps: (1) identification of studies through literature searches (studies repeated in the two searches were included only once); (2) screening for inclusion and exclusion criteria; (3) assessment for appropriate data extraction; (4) evaluation of study eligibility based on analysis conducted (excluding studies that did not report results from whole-brain analyses either in the main text or in the supplementary material); (5) attribution of the included studies to one or more contrasts (some of the studies reported coordinates contrasts that could be included in more than one contrast).

**Figure 2** shows the published activation foci from the included studies projected onto an inflated cortical surface for the contrast of a) VMI > Control, b) VMI > Perception, c) MMI > Control. See the supplementary videos on <u>Github</u> for a dynamical representation of the foci included in the VMI > Control, <u>VMI > Perception, MMI > Control</u> analyses.

**Figure 2.**
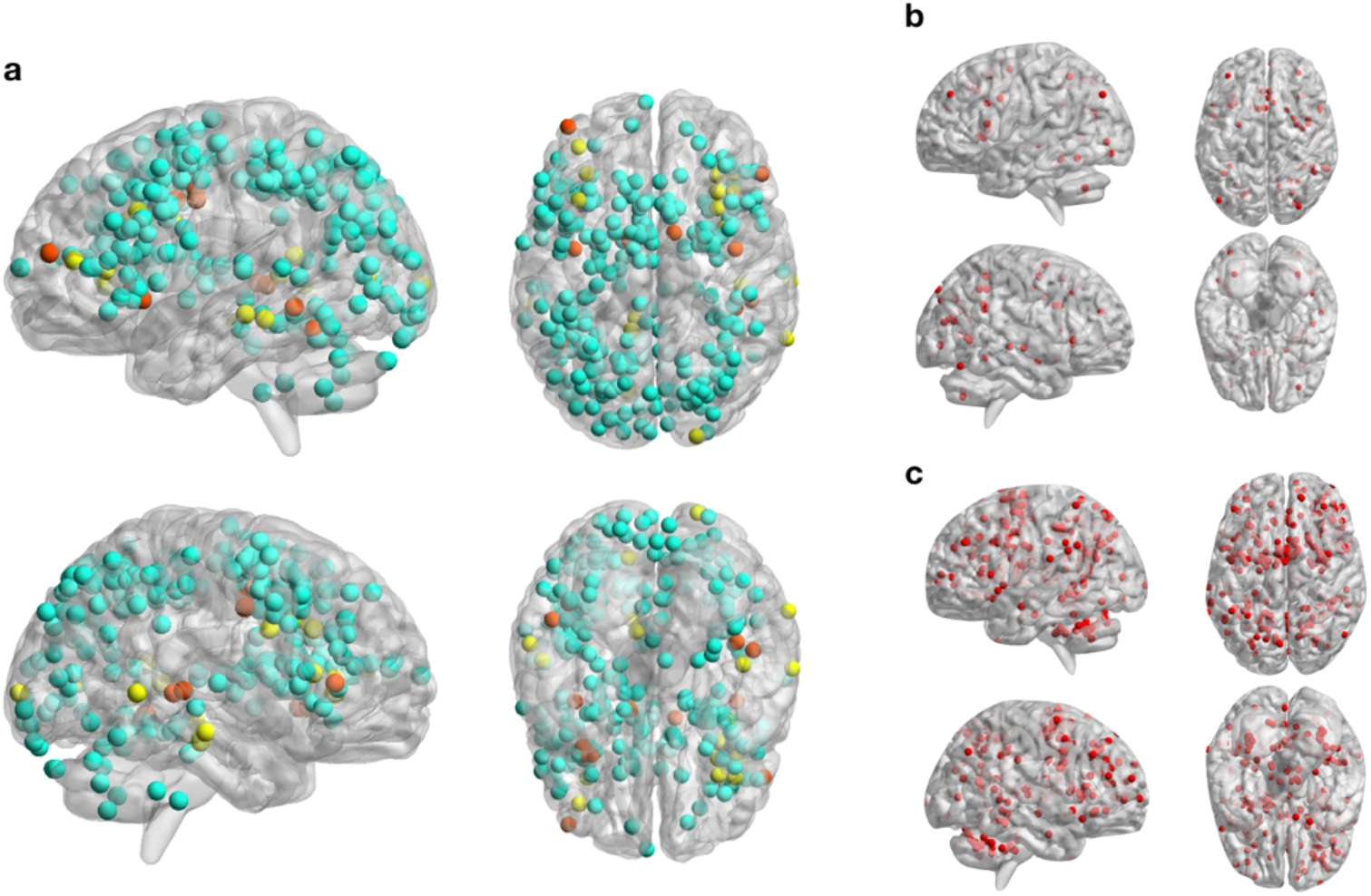
Activation foci reported in the studies included in the present meta-analysis for the contrasts a) VMI > Control, b) VMI > Perception, c) MMI > Control. For the VMI > Control panel, coordinates are color coded to represent the three categories of contrasts included: Object Form (cyan); Faces (yellow); Orthographic Material (orange).

### Activation likelihood estimation

GingerALE calculates the overlap of activation distributions across studies by modeling the foci as 3D Gaussian distributions centered at the reported coordinates with the full-width half-maximum (FWHM) weighted by the sample size of each study and combining the probabilities of activation for each voxel (Eickhoff et al., 2012). **Tables 1–3** report author names, publication year, type of stimuli used in the study, type of contrast, and number of subjects that were included in the contrasts VMI *versus* Control, VMI *versus* Perception, and MMI *versus* Control, respectively. Further, we conducted *conjunction* and *disjunction* analyses between (VMI *versus* Control) & (VMI *versus* Perception) as well as *conjunction* and *disjunction* analyses between (VMI *versus* Control) & (MMI *versus* Control).

**Table 1.**
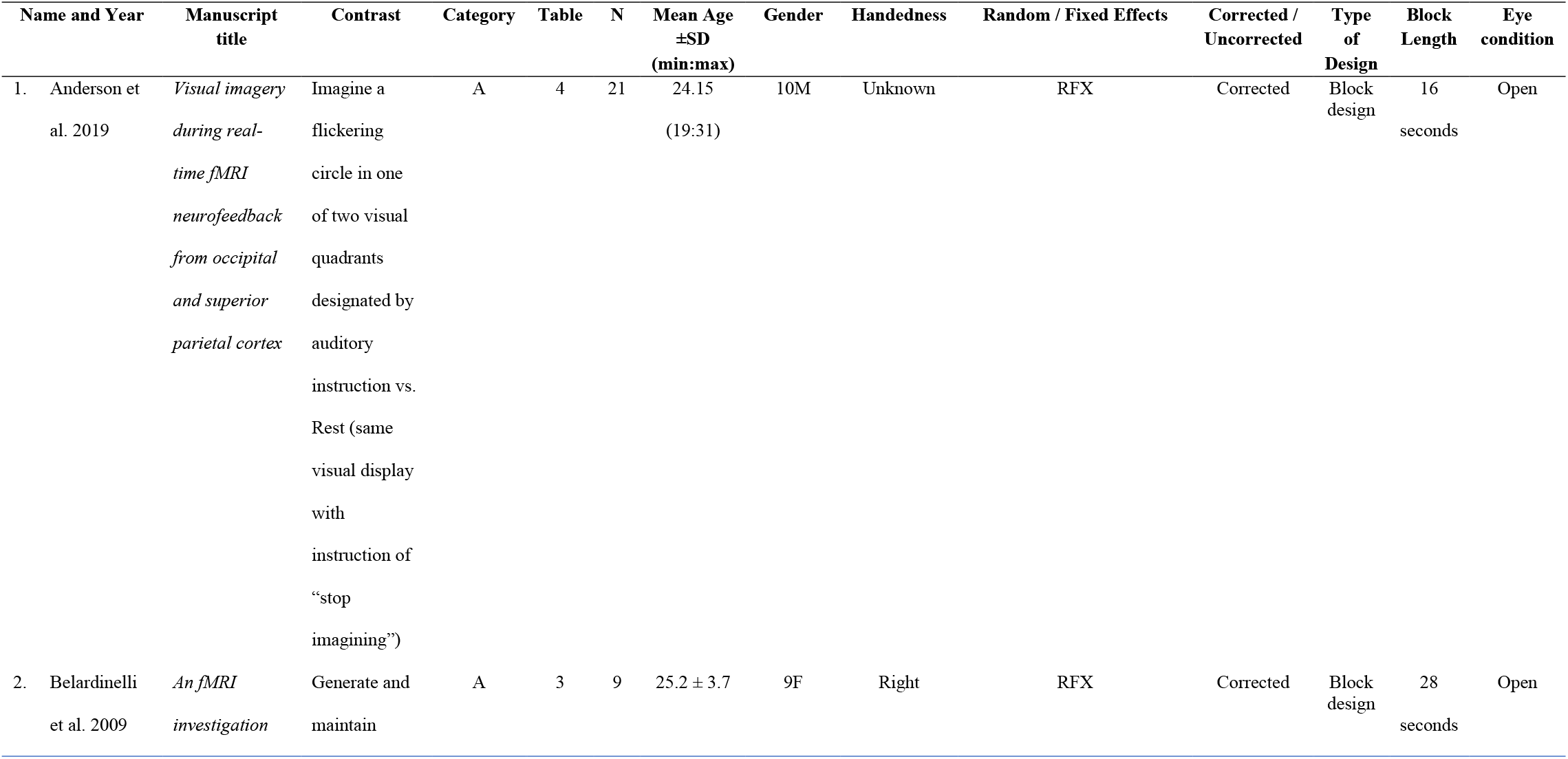
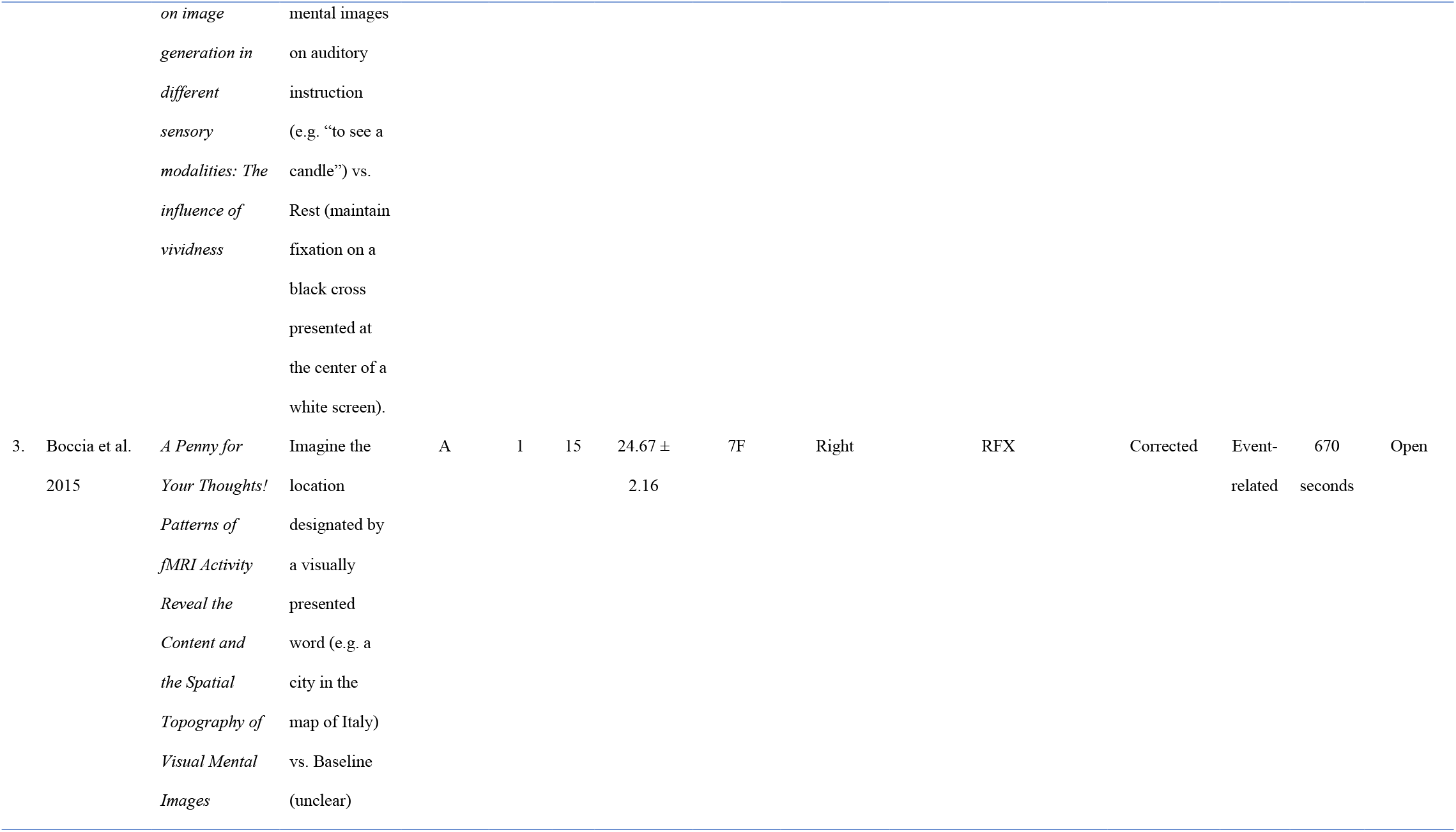
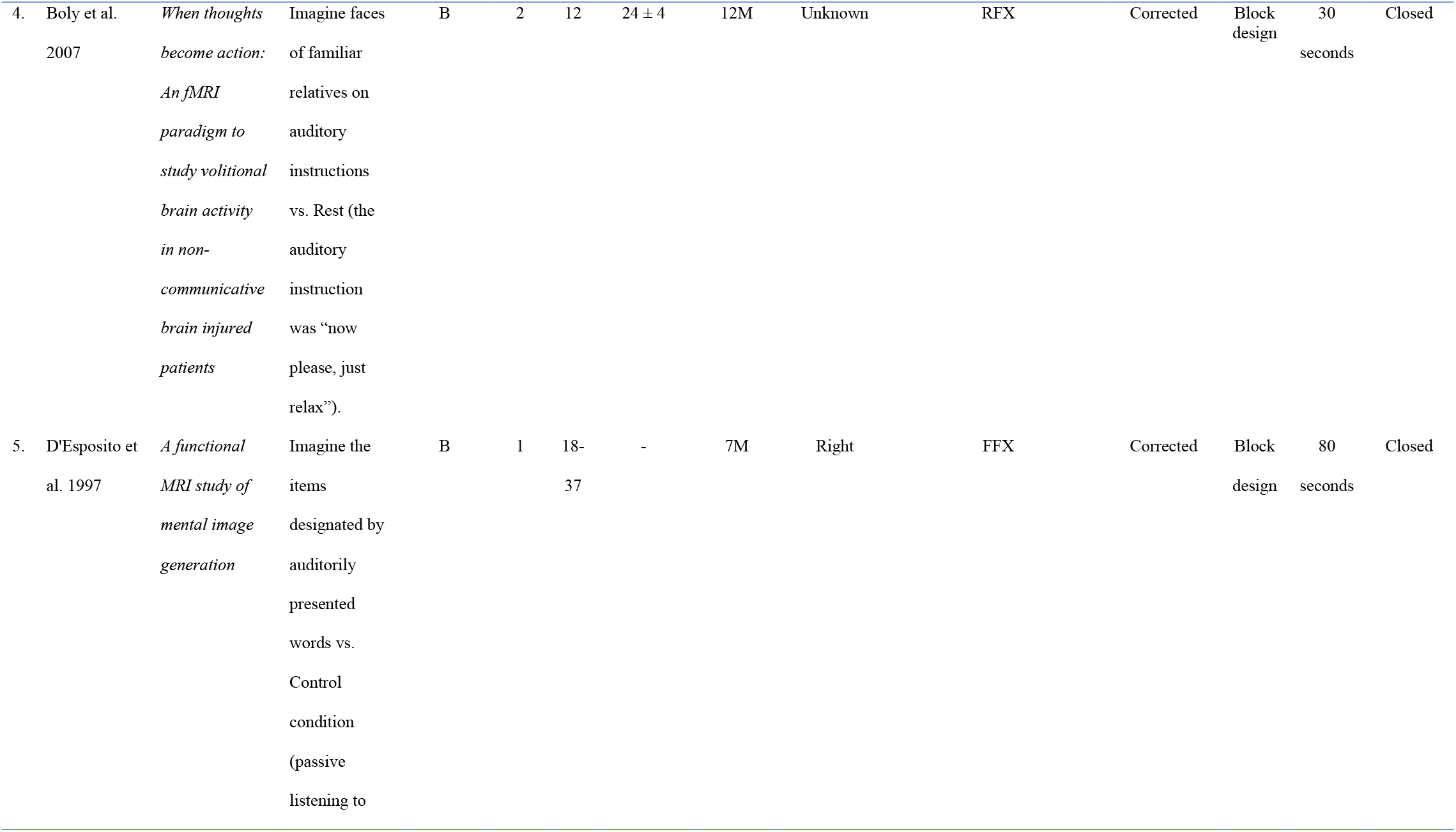
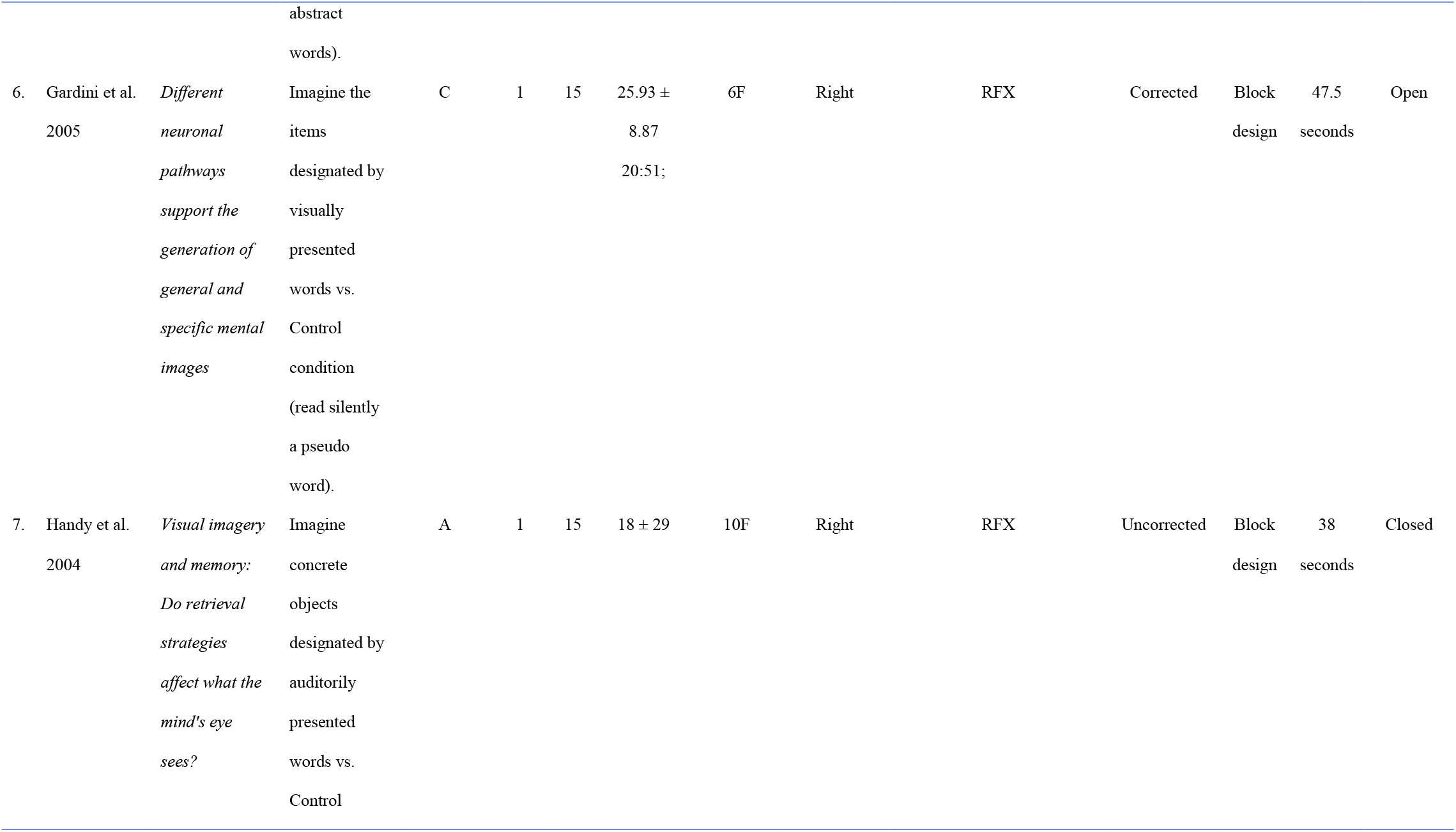
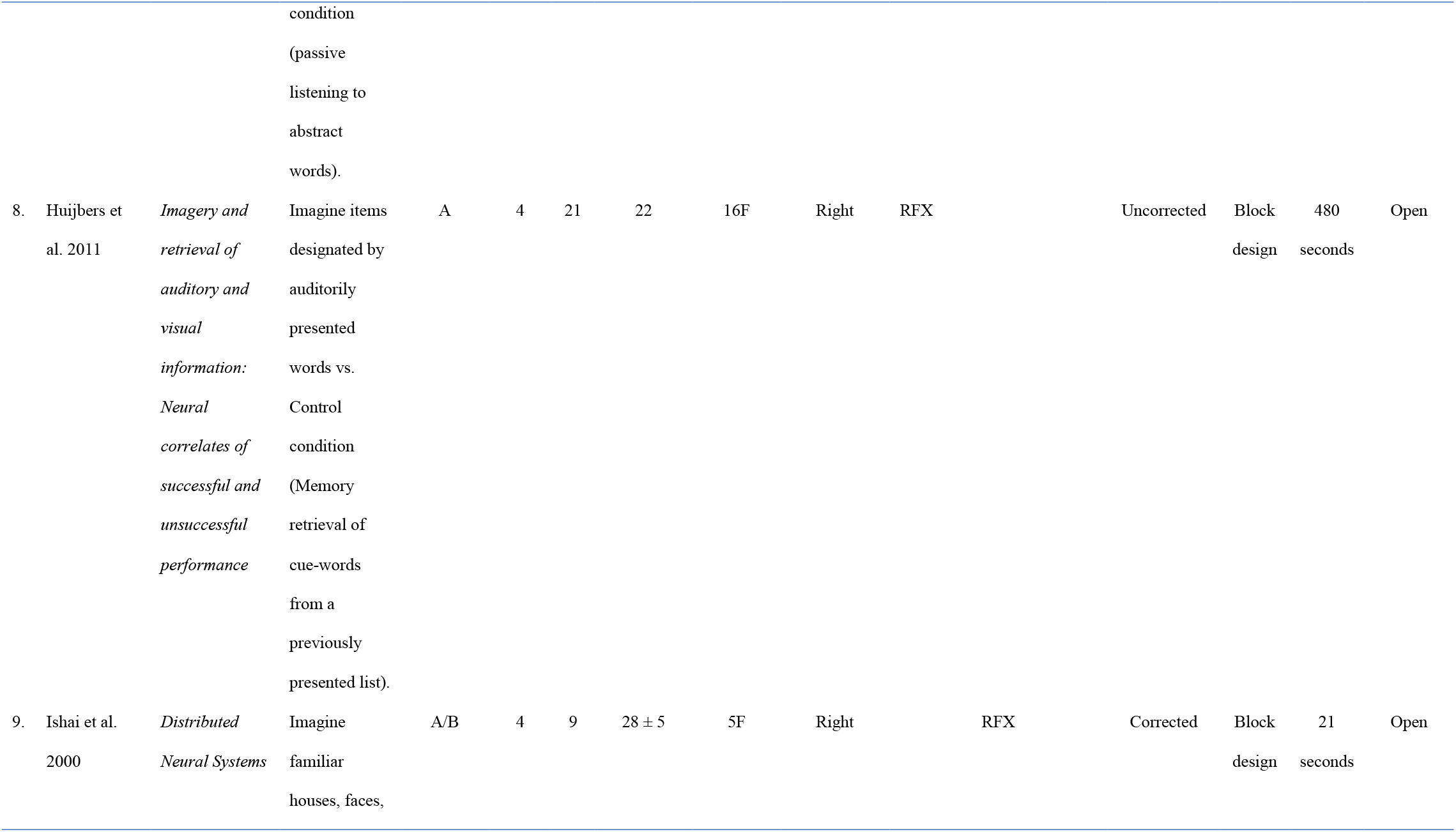
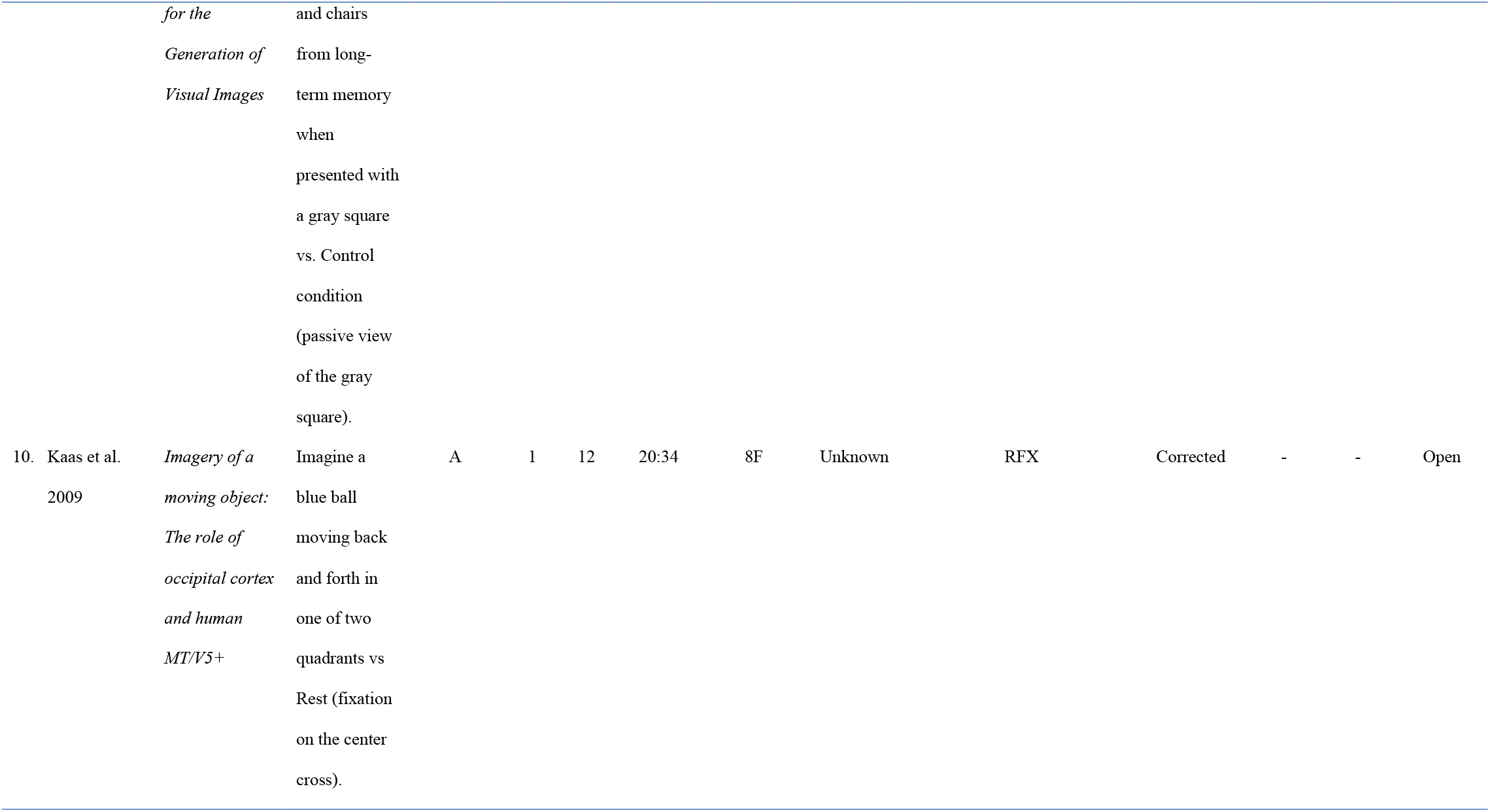
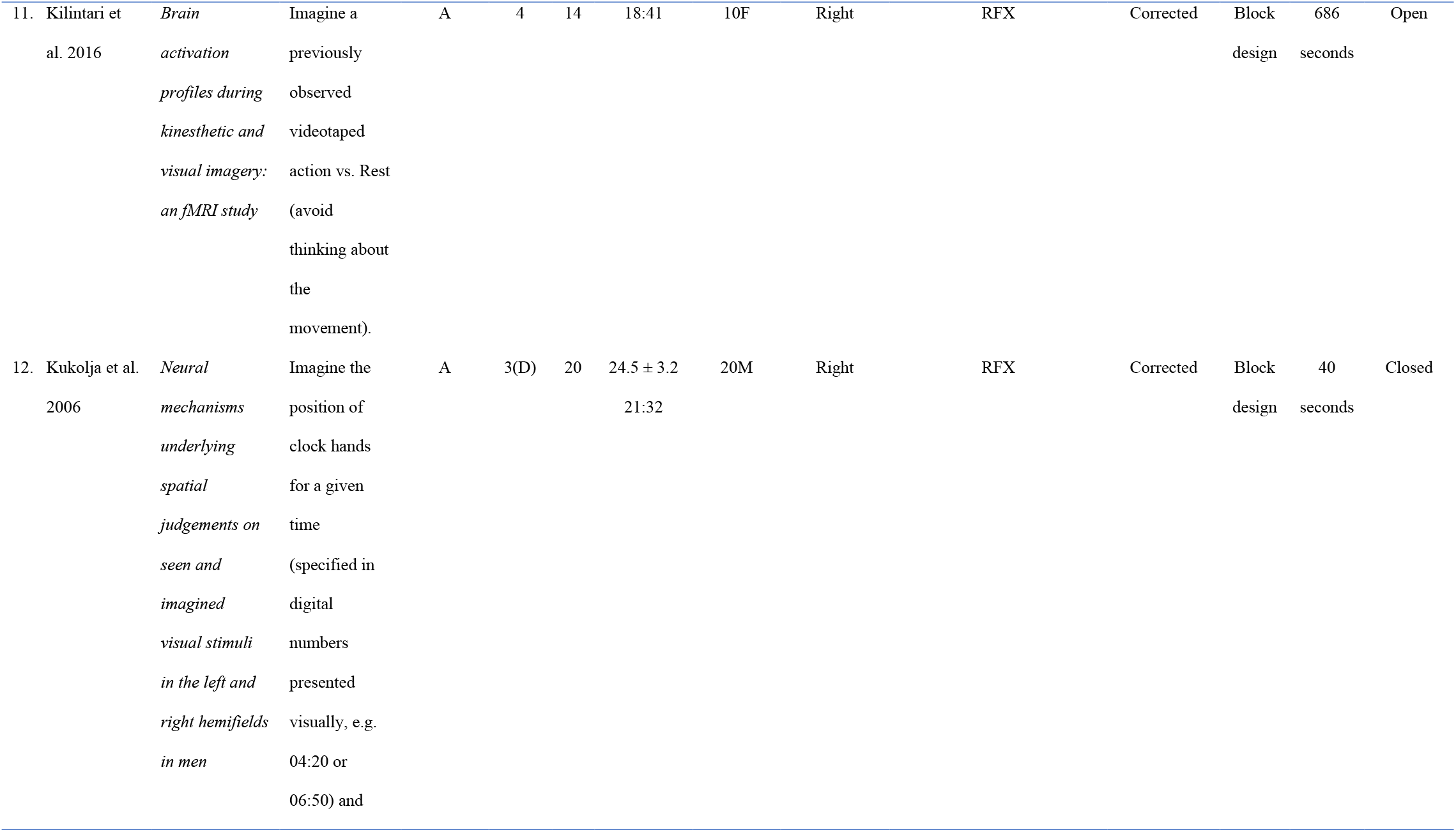
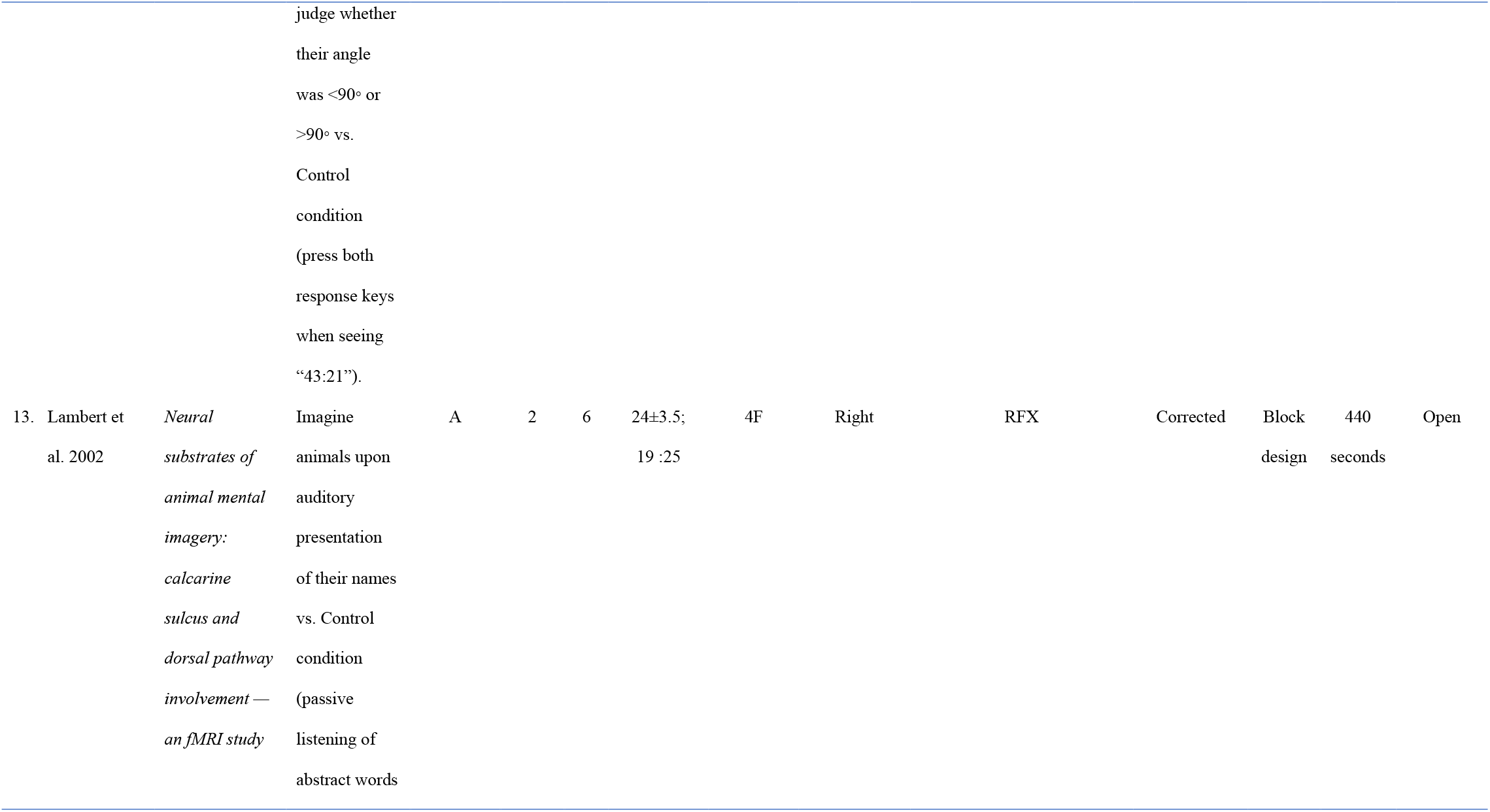
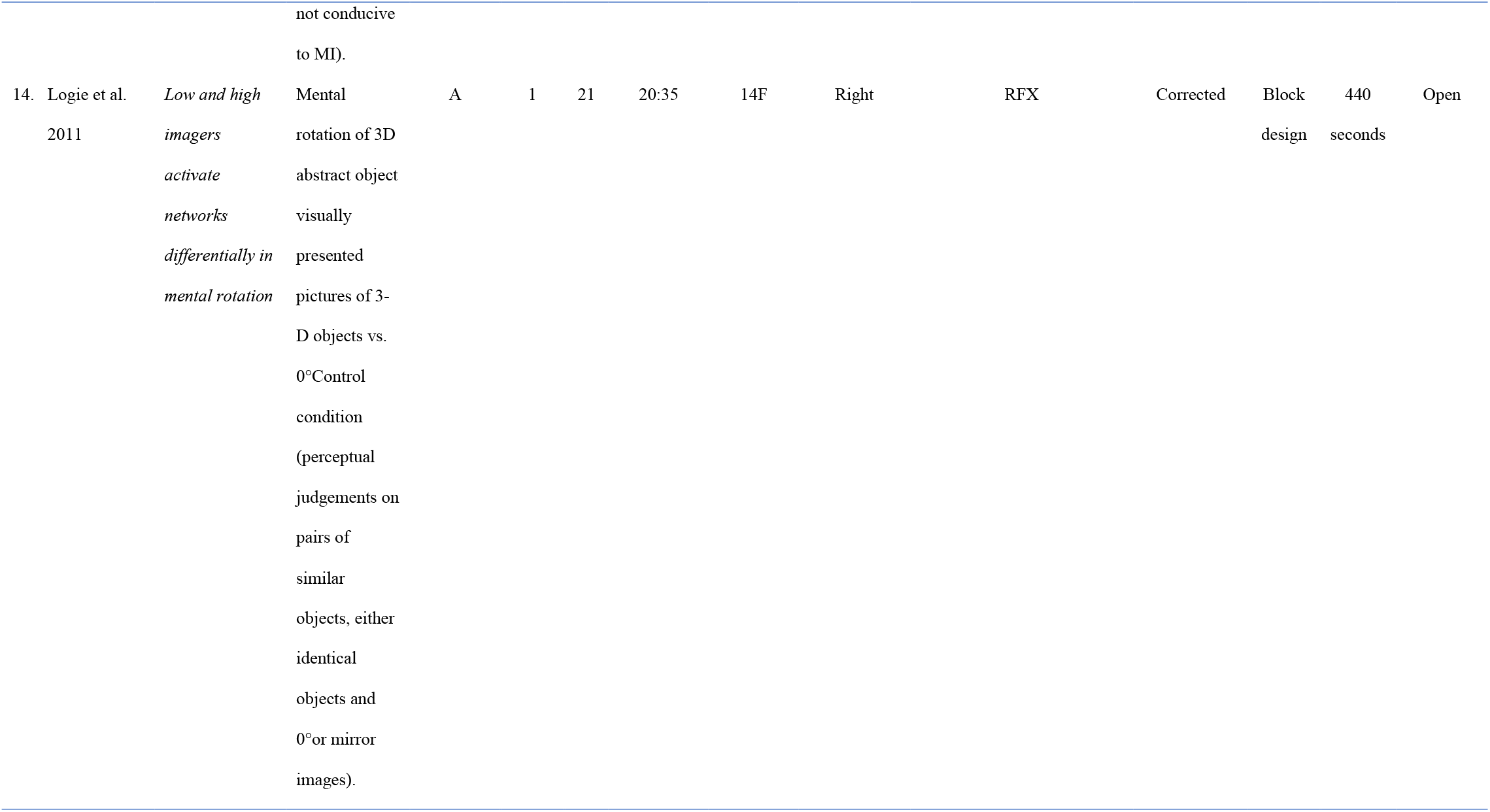
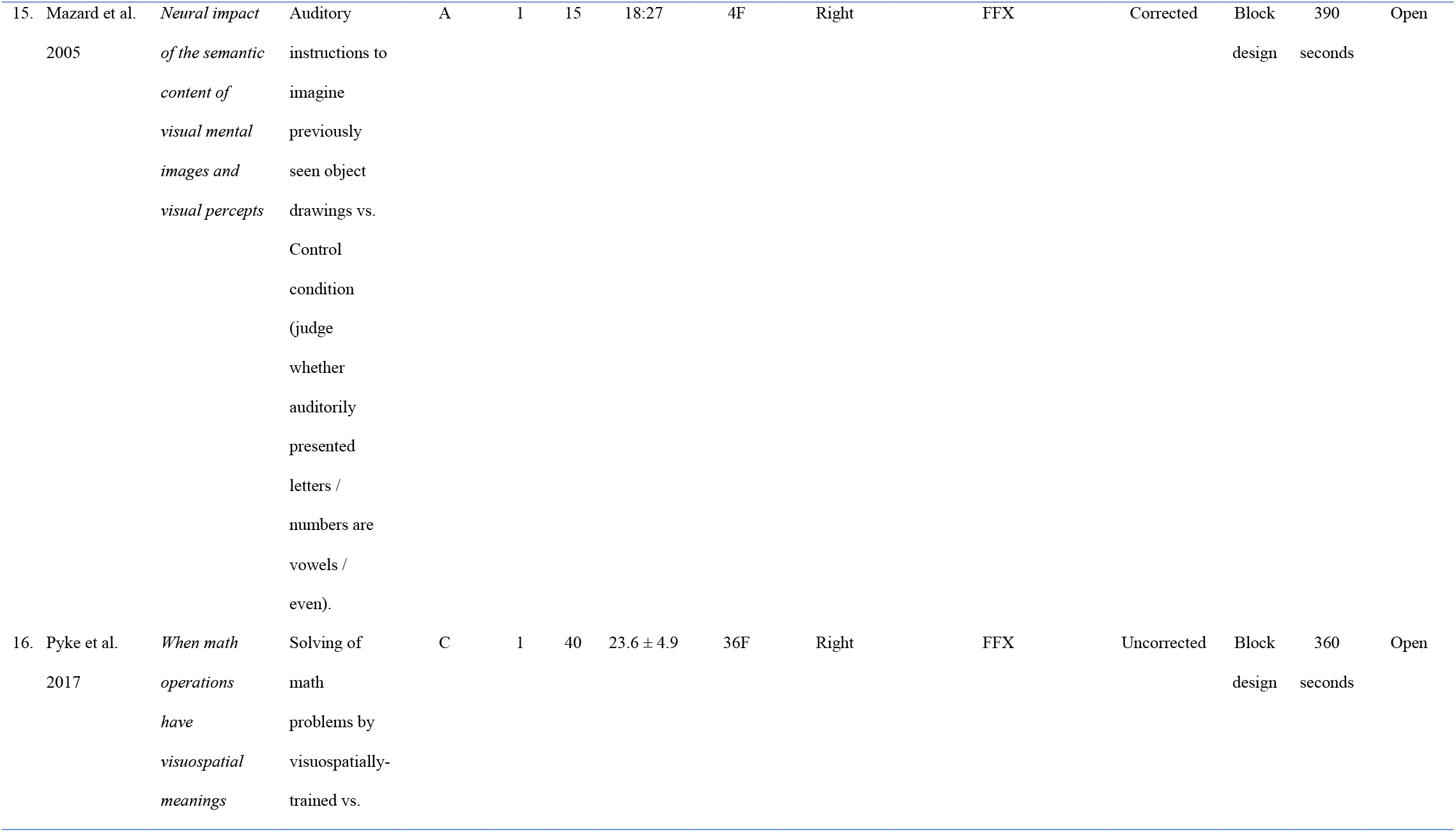
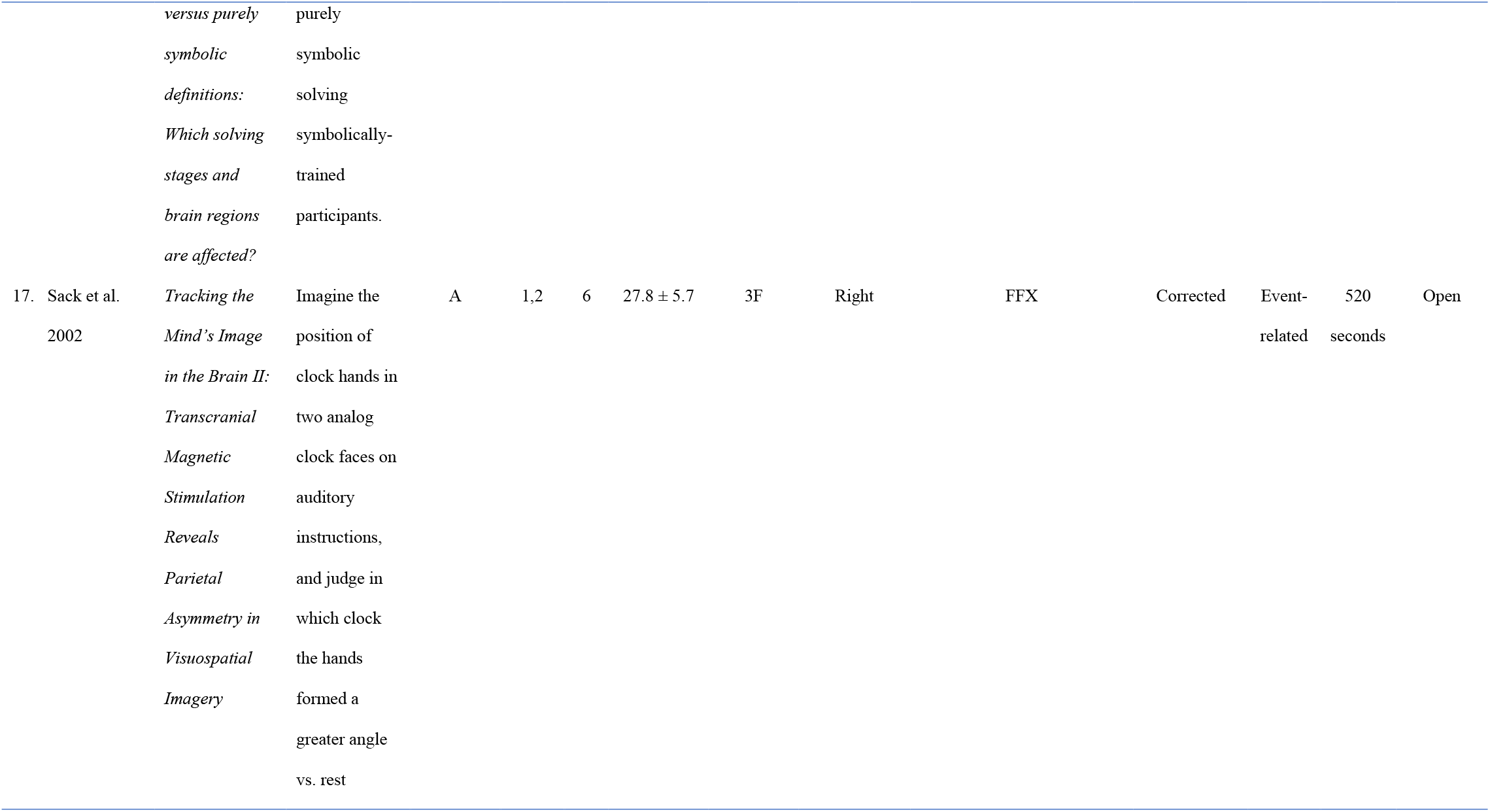
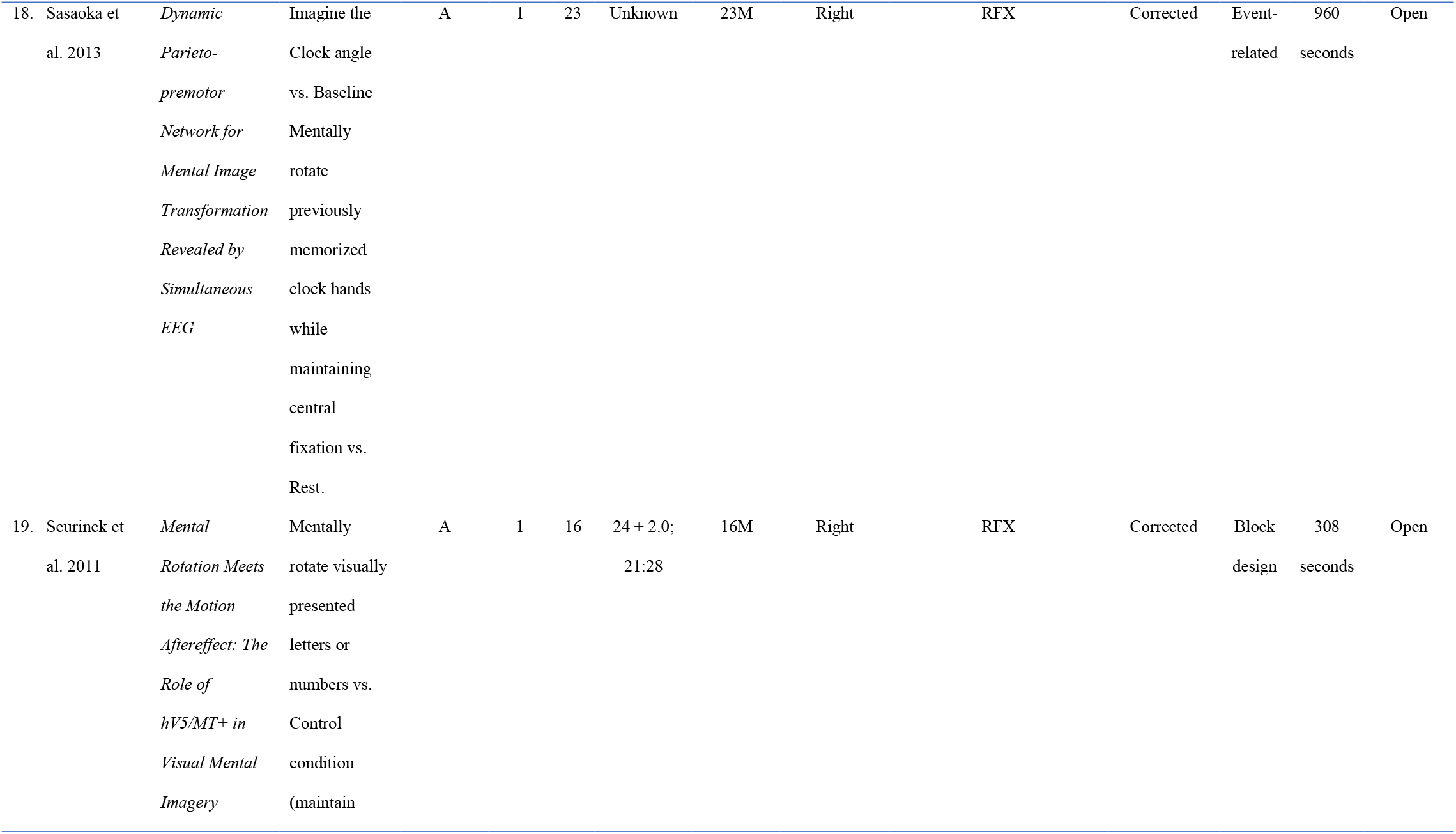
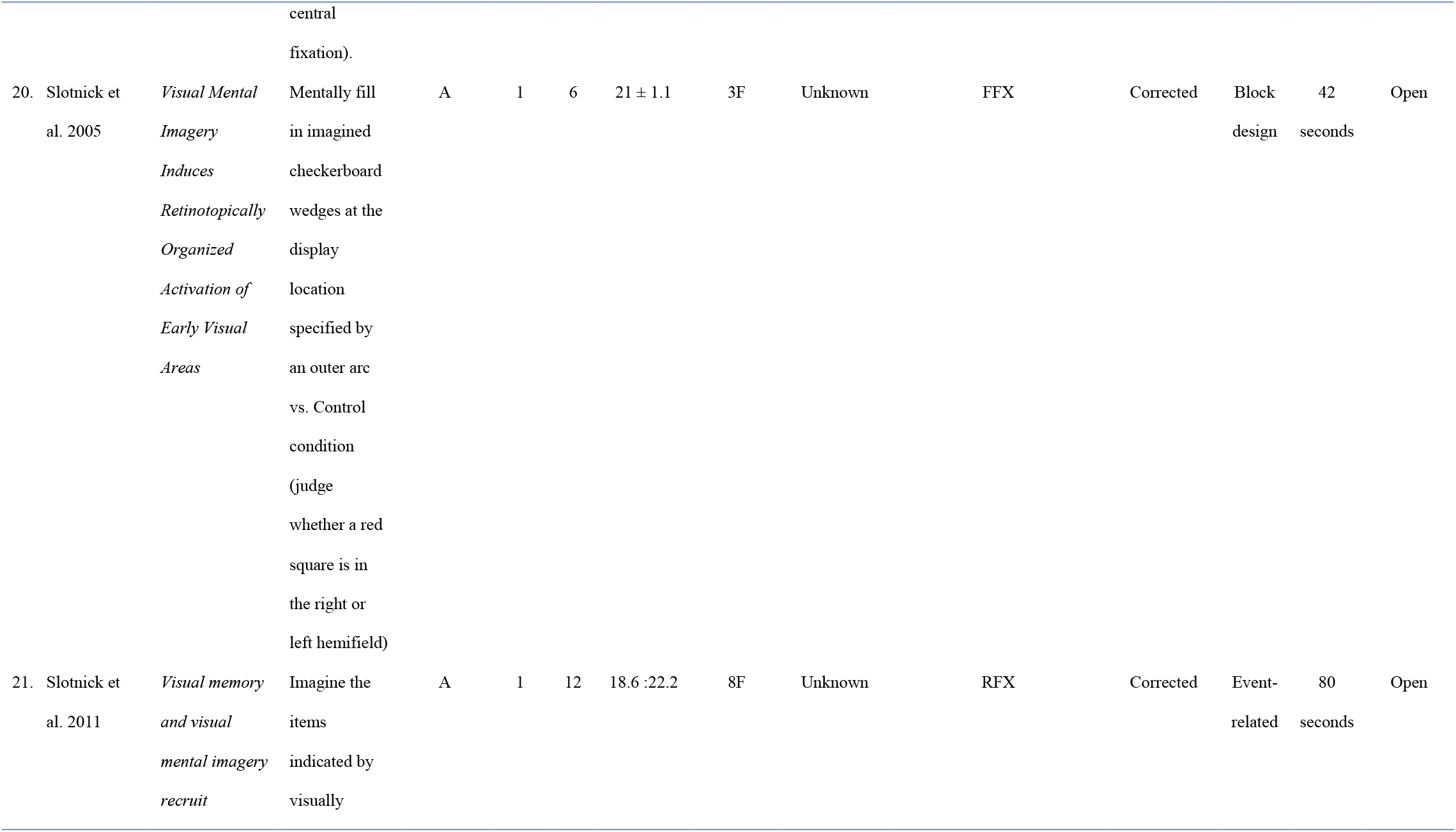
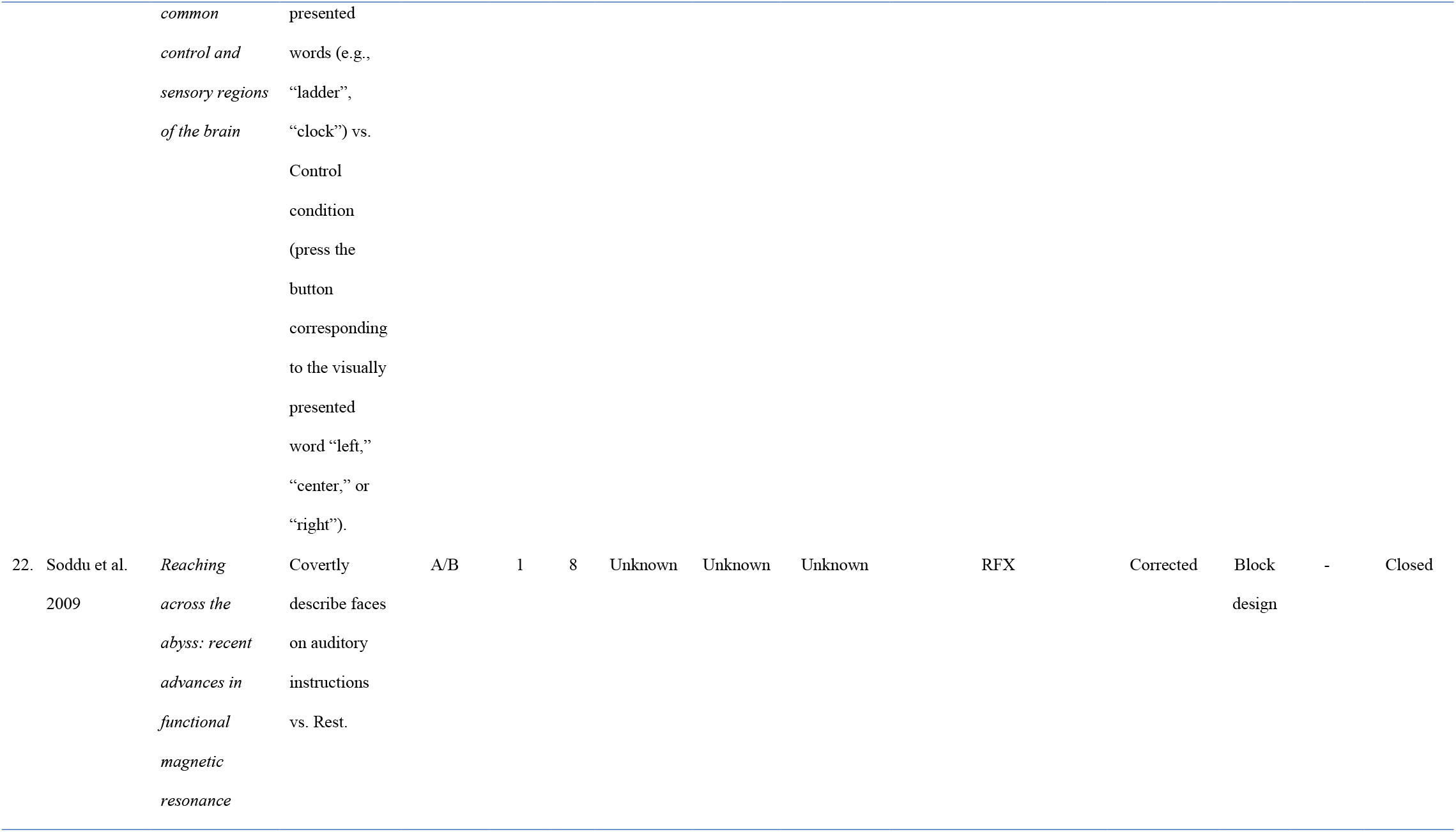
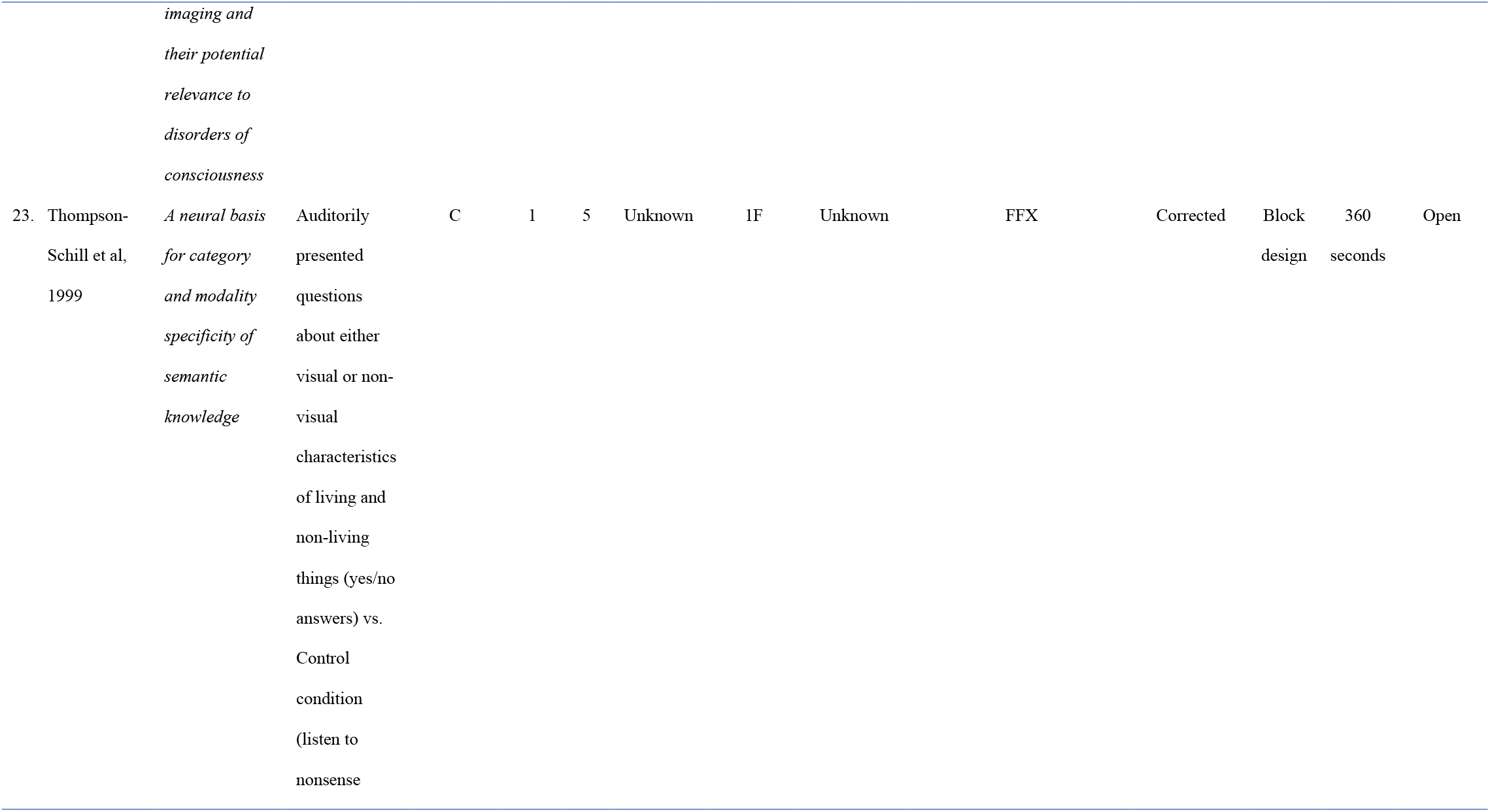
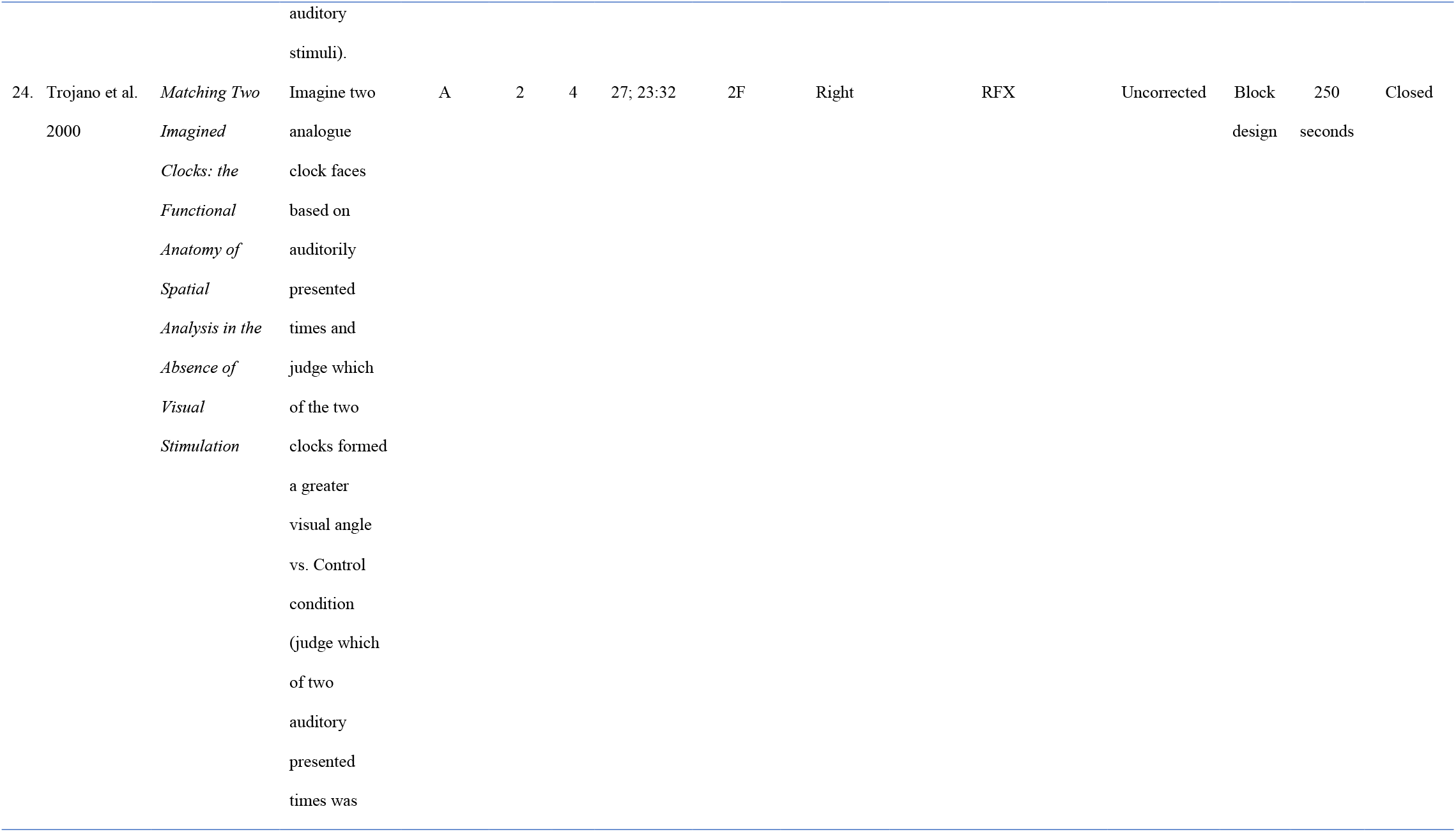
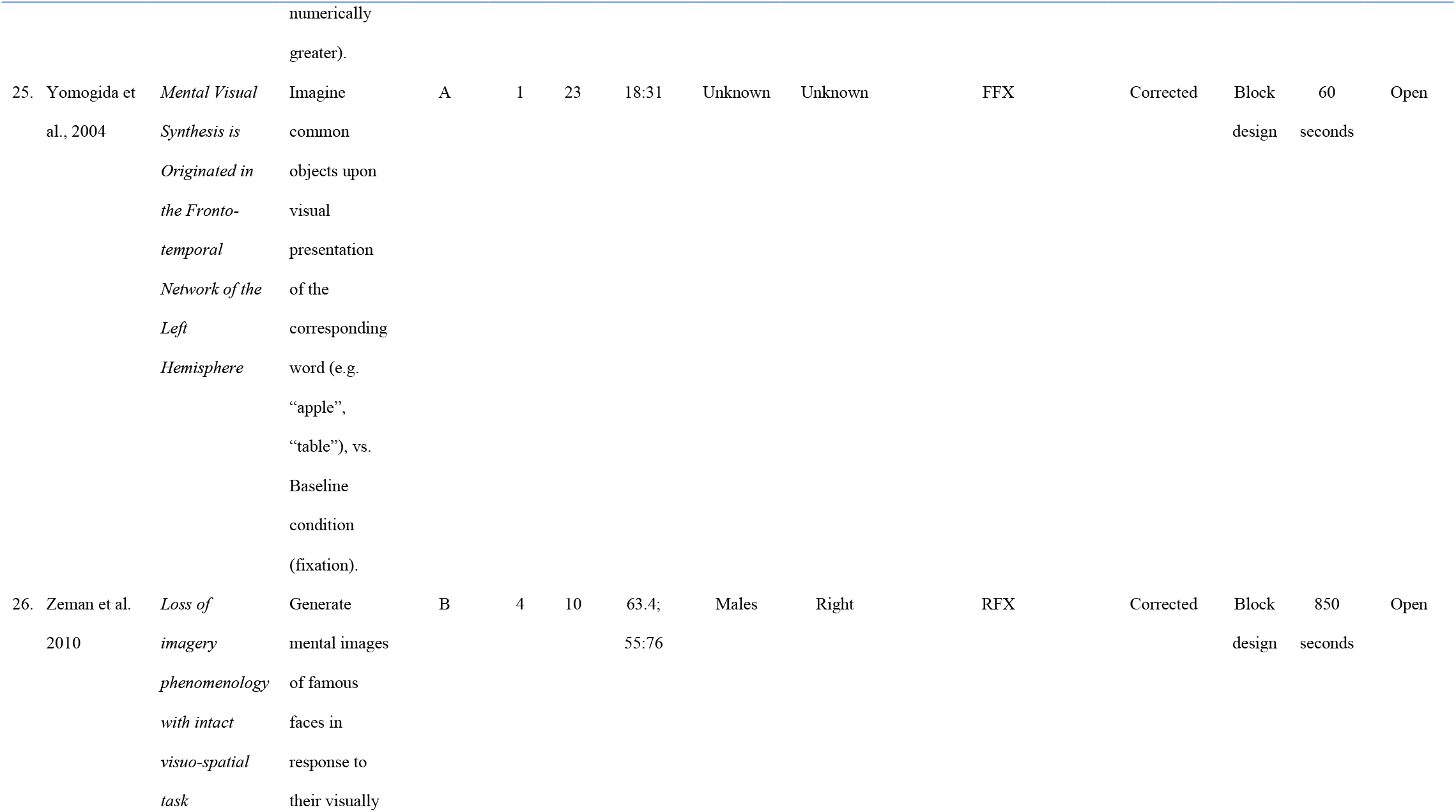
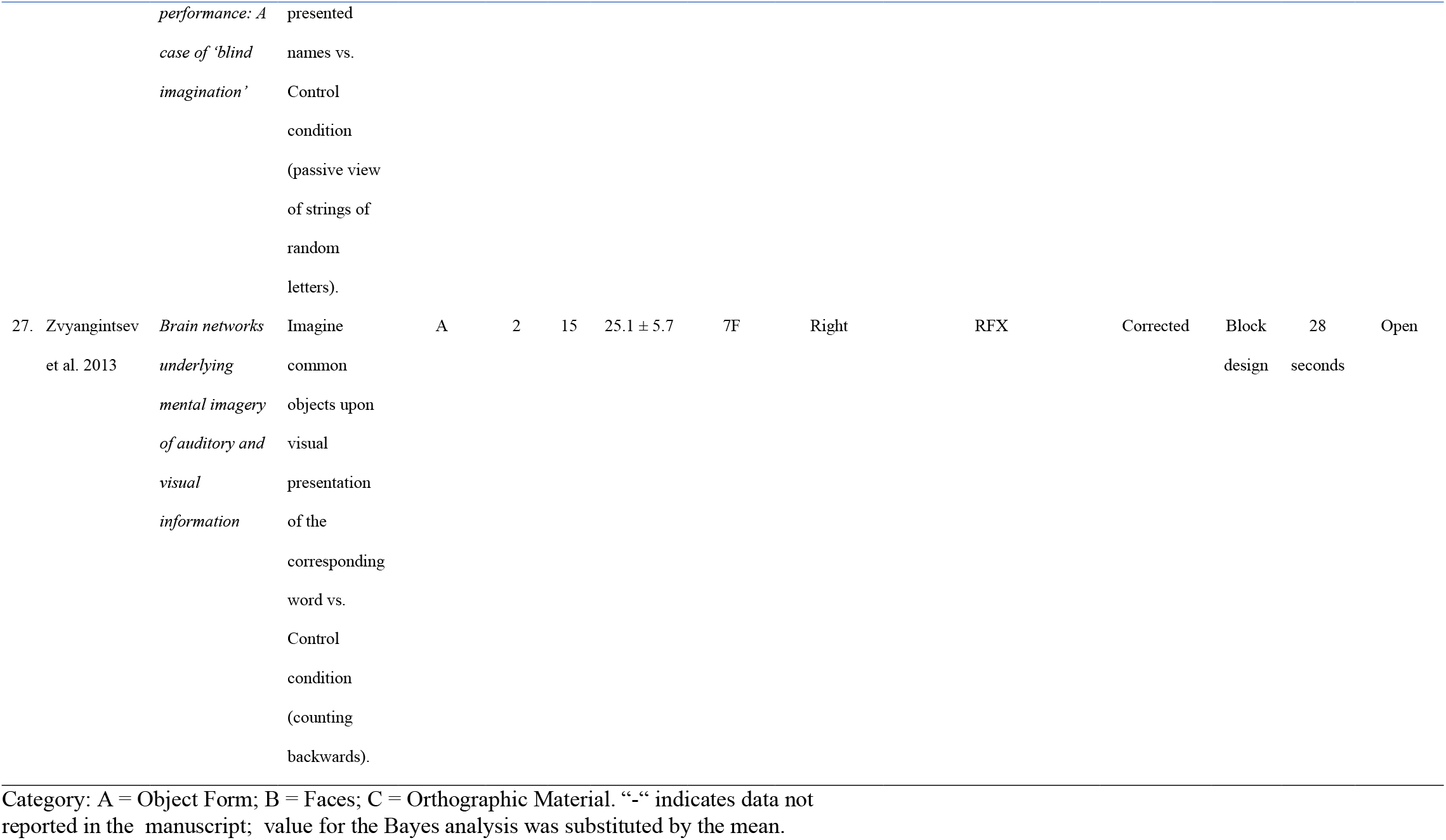
List of studies (n = 27) showing the first author’s name, publication year, manuscript title, contrast used, and number of subjects (n = 340) that were included in the ALE meta-analysis for the contrast Visual Mental Imagery > Control condition. Fixed or Random Effects and Corrected or Uncorrected variables were used as covariates in the Bayesian Analysis. A total of 372 activation foci were included in the analysis for this contrast.

Foci included in the dataset were either reported in MNI space or converted from Talairach space to MNI space by using the icbm2tal transform in GingerALE. The ALE map was assessed against a null-distribution of random spatial association across studies using a nonlinear histogram integration algorithm (Eickhoff et al., 2009; Eickhoff et al., 2012). All ALE maps were thresholded at a cluster-level corrected *p* < .05, with a cluster-forming *p* < .001 with 10,000 permutations for multiple comparison correction. Conjunction and disjunction analyses were then conducted at a *p* <.05 threshold and 10,000 permutations.

### Control analysis

A potential concern is that some of the control tasks used in fMRI experiments might in fact have engaged visual mental imagery (for example “read silently a pseudoword”, “perceptual judgements on pairs of similar objects”), contrary to the experimenters’ intention. To control for this possibility, we performed a follow-up analysis on studies that reported contrasts about visual mental imagery > rest (n = 10; Sack et al., 2002; Yomogida et al., 2004; Boly et al., 2007; Belardinelli et al., 2009; Soddu et al., 2009; Kaas et al., 2010; Seurinck et al., 2011; Sasaoka et al., 2014; Kilintari et al., 2016; Andersson et al., 2019).

### Spatial Bayesian Latent Factor Regression for Coordinate-Based Meta-Analysis

A specific aim of this study was to assess whether or not early visual activation occurred in the VMI vs Control condition, as a way to assess the potential contribution of early visual areas to VMI. Because the logic of our argument rests on testing the *absence* of activation, we conducted a spatial Bayesian latent factor regression model, where the probability that a voxel is reported as an activated focus, is modeled via a doubly stochastic Poisson process (Montagna et al., 2018). To test the activation degree of each voxel, the null hypothesis (*H*_0_) stated that the voxel was specifically activated as a focus during VMI vs Control condition. We took the posterior probability *p*(*v*) as the probability of null hypotheses for each voxel. The alternative hypothesis (*H*_1_) was that the activation of the voxel belongs to the ongoing brain activity and was not specific. Thus, we took the mean activation probability of the whole brain voxels to get the H1 probability (see (Montagna et al., 2018) for additional details about the method and a general discussion of the properties of this model). We defined Bayes Factors (Rouder et al., 2009) for accepting and rejecting the null hypothesis. To compute the odds hypothesis conditionally on observed data *Bayes Factors = Pr(H0\data)/Pr(H1\data):* BF (odds) > 3 were considered as the criteria to accept null hypothesis. Bayes Factors (BF) (odds) < 0.33 were considered as the criteria to reject the null hypothesis and thus to accept the alternative hypothesis. Spatial Bayesian latent factor regression analysis was conducted on the dataset consisting of 27 studies on VMI > Control condition for a total of 376 foci (see **Table 1** for the list of studies included in this analysis). For each study, we also included five supplementary covariates: inference method (fixed / random effects), *p*-value correction (corrected / uncorrected), number of participants, type of design (block or event-related), block length (in seconds), and eye condition (closed or open). We used a standard brain mask of 91 x 109 x 91 voxels of 2mm in size. Kernels falling outside the mask were discarded. We assigned a Gamma(1, 0.3) prior distribution with mean 1/3 to the diagonal elements of *Σ*^-1^. The sampler was run for 10,000 iterations and the algorithm converged, with the first 5,000 samples discarded as a burn-in and collecting every 50th sample to thin the chain. We obtained the posterior intensity values of the full brain voxels, which we used to compute the BFs.

## Results

The meta-analysis conducted on all the coordinates from the 46 experiments (*all mental imagery*) showed activation increases in areas within the fronto-parietal networks (including the bilateral superior frontal gyrus, and the bilateral inferior parietal lobe), and the bilateral anterior insular cortices. In addition, unilateral activation increases consistent across the studies were found in the right precuneus and in the anterior portion of the left calcarine sulcus, in the left fusiform and parahippocampal gyri. **Figure 3** shows the section views of the brain and highlights the regions with consistent activation increase across all domains, while **Table 4** reports the coordinates of common activation across all domains.

**Figure 3.**
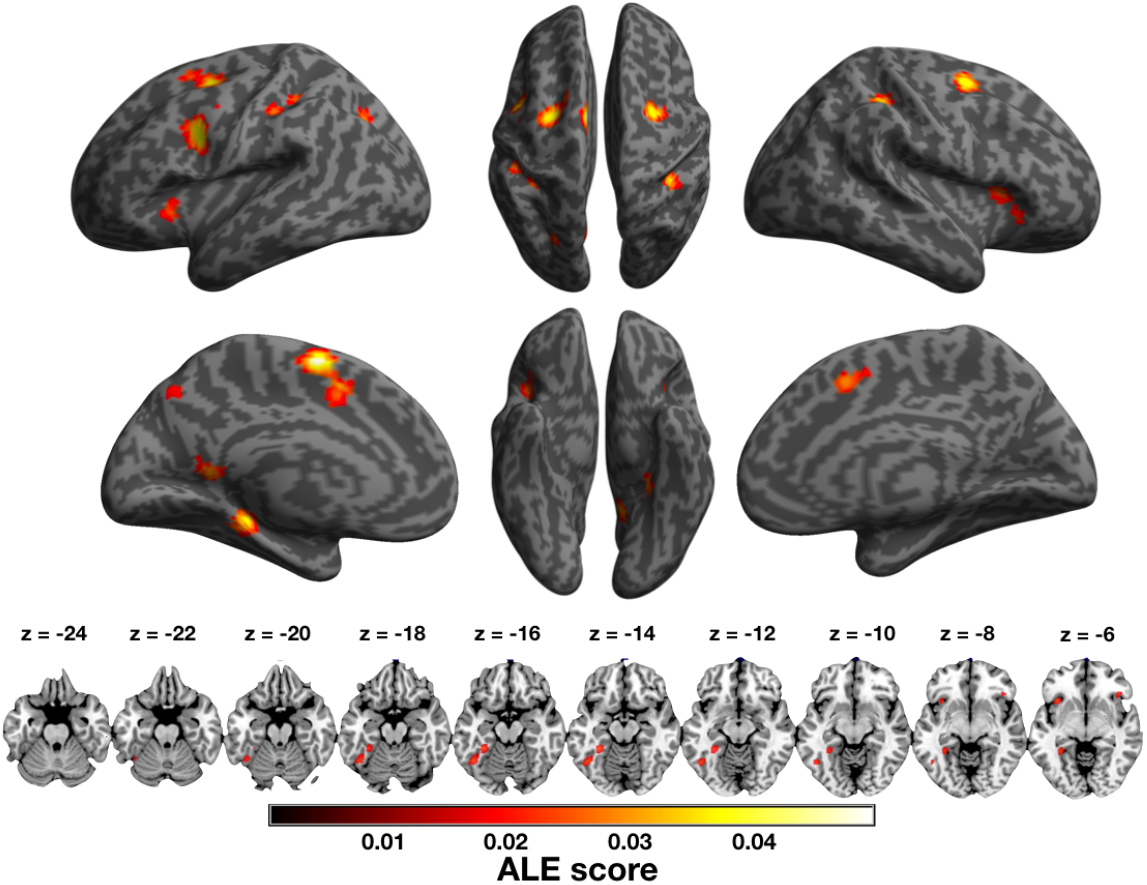
Brain regions showing consistently increased activation associated with mental imagery (across all the domains of VMI > Control, VMI > Perception, and MMI > Control).

**Table 2.** List of studies (n = 4) showing the first author’s name, publication year, manuscript title, contrast used, and number of subjects (n = 52) that were included in the ALE meta-analysis for the contrast Visual Mental Imagery > Perception condition. Fixed or Random Effects and Corrected or Uncorrected variables were used as covariates in the Bayesian Analysis. A total of 62 activation foci were included in the analysis for this contrast.

| Name and Year | Manuscript title | Contrast | Table | N | Mean Age ±SD<br>(min:max) | Gender | Handedness | Random /<br>Fixed<br>Effects | Corrected /<br>Uncorrected |
| --- | --- | --- | --- | --- | --- | --- | --- | --- | --- |
| 1. Arzy et al.<br>2009 | *Subjective mental time: the functional architecture of projecting the self to past and future* | Mental time imagery - self projection in different time points and events vs. Face discrimination | 1 | 12 | 33.7 ± 4.2<br>(29:38) | 5F | Right | FFX | Corrected |
| 2. Daselaar et al. 2010 | *Modality-specific and modality-independent components of the human imagery system* | Visual Imagery vs. Visual Perception | 2 | 16 | 21 | 10F | Right | RFX | Uncorrected |
| 3. Kukolia et al. 2006 | *Neural mechanisms underlying spatial judgements on seen and imagined visual stimuli in the left and right hemifields in men* | Visual Imagery vs. Visual Perception | 3 | 20 | 24.5 ± 3.2<br>(21:32) | 20M | Right | RFX | Corrected |
| 4. Trojano et al. 2000 | *Matching Two Imagined Clocks: the Functional Anatomy of Spatial Analysis in the Absence of Visual Stimulation* | Visual imagery vs. spatial matching of visually presented clocks | 3 | 4 | 27; (23:32) | 2F | Right | FFX | Uncorrected |

**Table 3.**
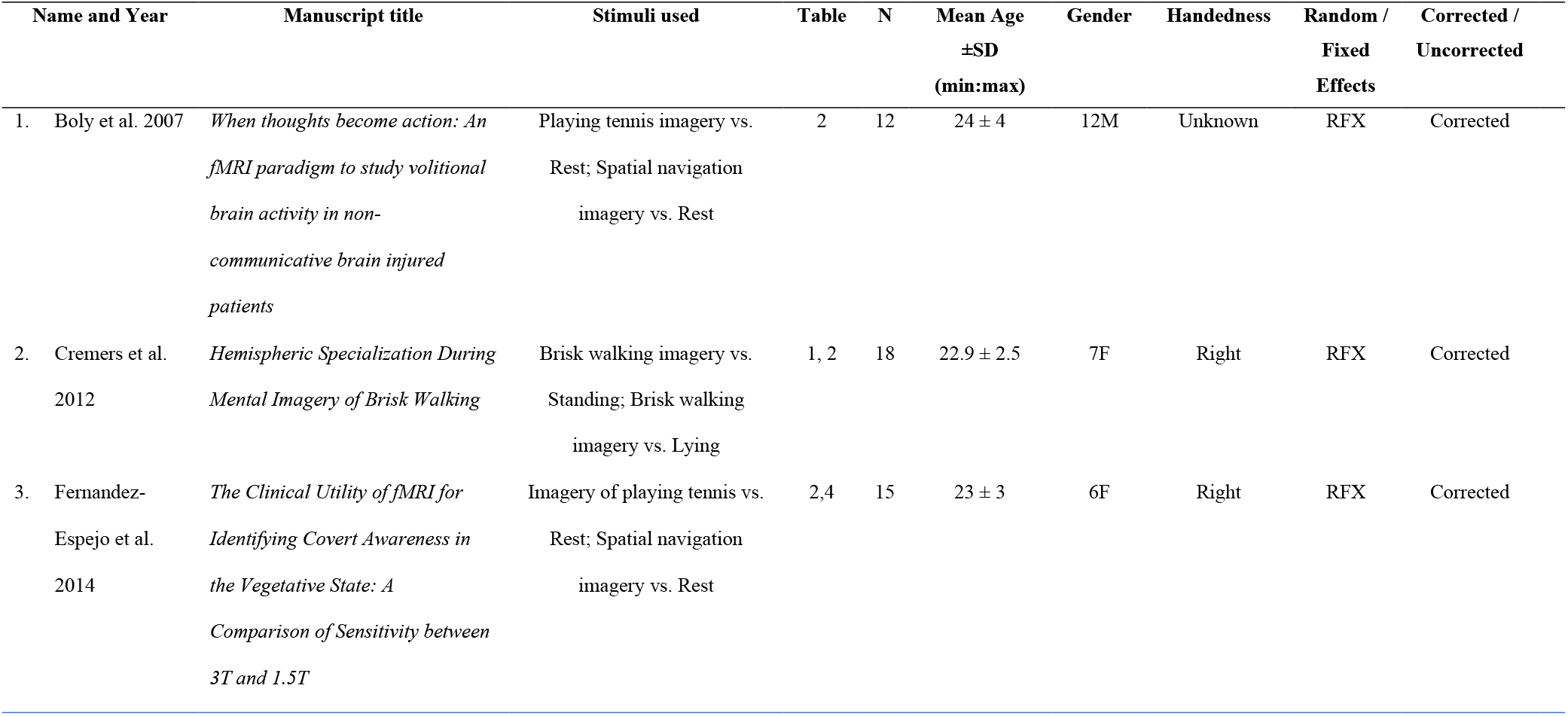
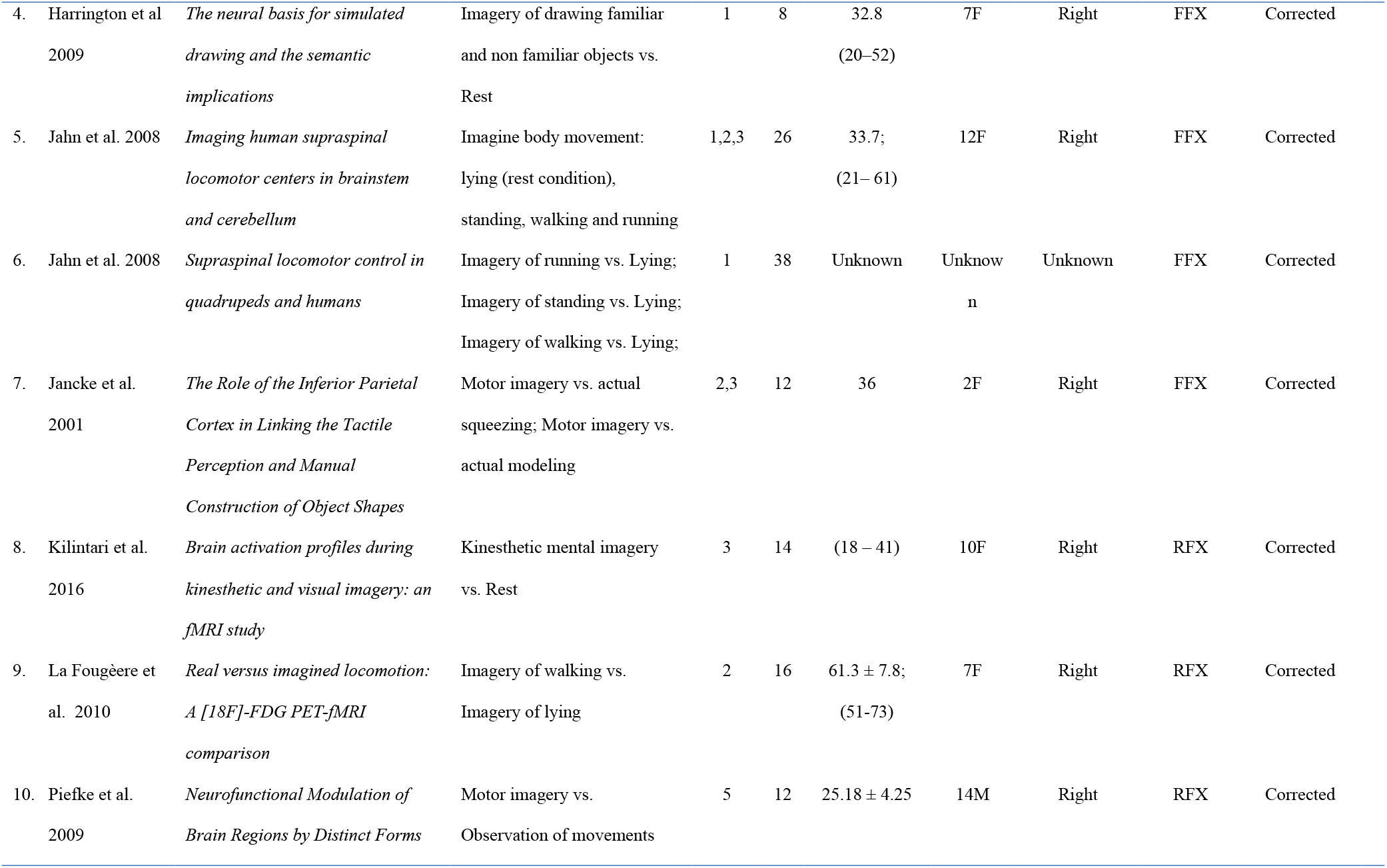
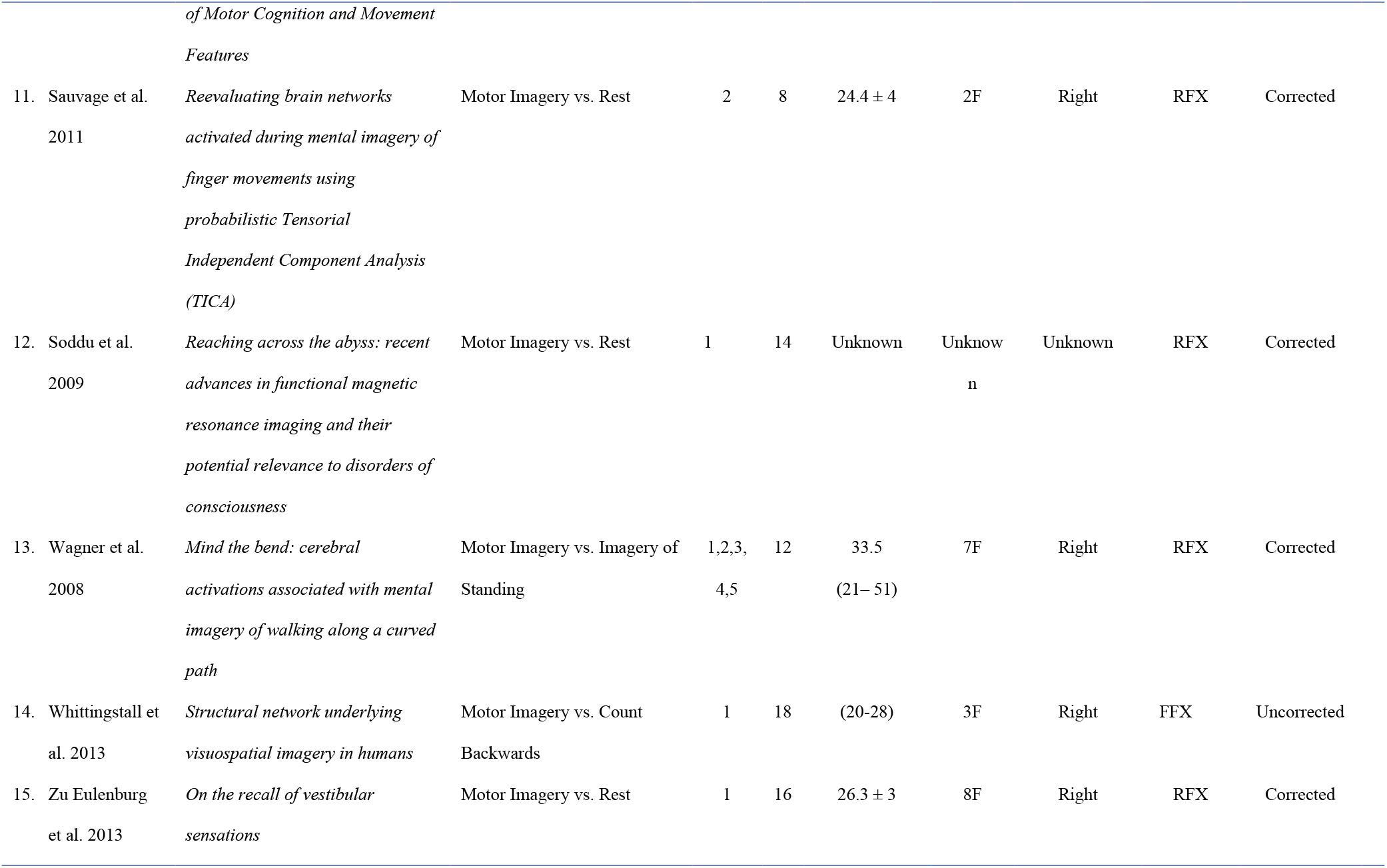
List of studies (n = 16) showing the first author’s name, publication year, manuscript title, stimuli used, and number of subjects (n = 282) that were included in the ALE meta-analysis for the contrast Motor Mental Imagery > Control condition. Fixed or Random Effects and Corrected or Uncorrected variables were used in the Bayesian Analysis. A total of 346 activation foci were included in the analysis for this contrast.

**Table 4.** Brain regions showed consistent increased activation across all the mental imagery domains (46 studies, 787 foci, 671 subjects).

| Region | L/R | BA | x | y | z | Z | K |
| --- | --- | --- | --- | --- | --- | --- | --- |
| Supp. Motor Area | L | 6 | -4 | -2 | 64 | 0.05 | 1020 |
| Precentral Gyrus<sup>a</sup> | R | 6 | 30 | -4 | 54 | 0.04 | 463 |
| Insular Cortex | L | 47 | -30 | 22 | 4 | 0.04 | 336 |
| Precentral Gyrus<sup>a</sup> | L | 6 | -28 | -4 | 56 | 0.04 | 511 |
| Inferior Parietal Lobe | L | 40 | -38 | -42 | 46 | 0.04 | 503 |
| Insular Cortex | R | 47 | 34 | 22 | 2 | 0.03 | 357 |
| Inferior Parietal Lobe | R | 40 | 40 | -42 | 48 | 0.03 | 297 |
| Middle Occipital Gyrus | L | 19 | -26 | -76 | 38 | 0.03 | 741 |
| Cerebellum Crus1 | L | 0 | -32 | -60 | -36 | 0.03 | 189 |
| Precentral Gyrus | L | 6 | -52 | 6 | 36 | 0.03 | 436 |
| Precuneus | R | 7 | 16 | -68 | 50 | 0.03 | 114 |
| Calcarine Sulcus | L | 17 | -12 | -56 | 10 | 0.03 | 266 |
| Fusiform Gyrus | L | 37 | -46 | -58 | -14 | 0.02 | 149 |
| Parahippocampal Gyrus | L | 37 | -32 | -44 | -4 | 0.02 | 161 |

**Note:** x, y, z: coordinates of the corresponding peak in MNI space, in units of millimeter.
Abbreviations: ALE, activation likelihood; BA, Brodmann area; K, cluster size in number of voxels; L, left; R, right. <sup>a</sup> comprises the Frontal Eye Fields.

The meta-analysis of the 27 experiments included in the VMI *vs* Control condition showed activation increases in bilateral superior frontal gyri, in the left anterior insular cortex, in a large cluster with a peak in the bilateral inferior parietal lobe extending towards the superior parietal lobe and to the superior portion of the middle occipital gyrus (BA19), in the left middle frontal gyrus, in the left fusiform gyrus, as well as regions lateralized to the right hemisphere, including the angular gyrus, the supplementary motor area / anterior cingulate cortex (**Figure 4** and **Table 5**).

**Figure 4.**
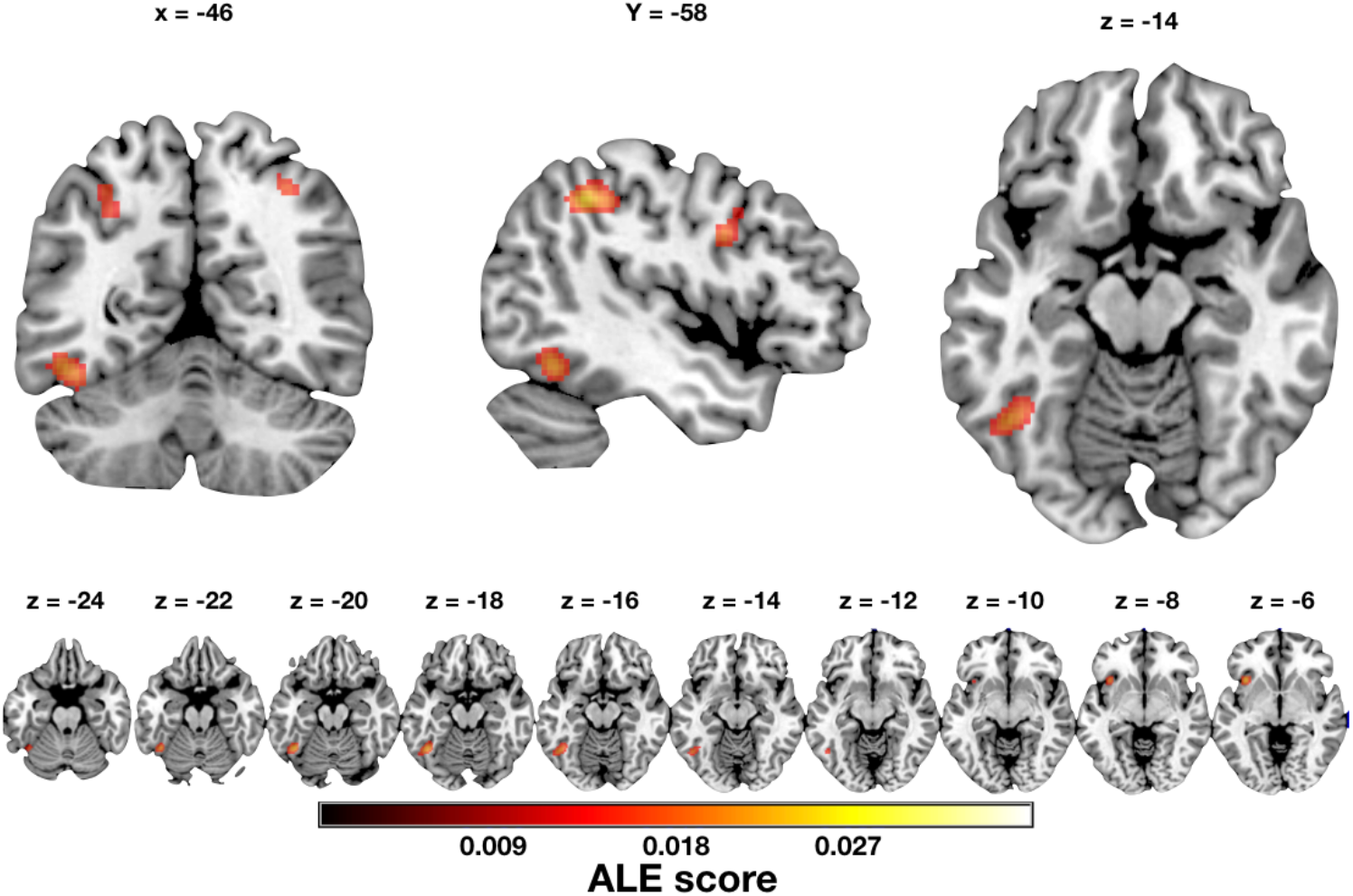
Brain regions showing consistently increased activation associated with the contrast VMI > Control condition, including our hotspot in the left fusiform gyrus (which we labeled the Fusiform Imagery Node, see Discussion), and fronto-parietal areas bilaterally.

**Table 5.** Brain regions showed consistent increased activation in the VMI *versus* Control condition (27 studies, 376 foci, 380 subjects).

| Region | L/R | BA | x | y | z | Z | K |
| --- | --- | --- | --- | --- | --- | --- | --- |
| Precentral Gyrus | R | 6 | 30 | -4 | 54 | 0.03 | 270 |
| Inferior Parietal Lobe | L | 40 | -40 | -44 | 46 | 0.03 | 910 |
| Superior Parietal Lobe | L | 7 | -16 | -64 | 52 | 0.02 |  |
| Middle Occipital Gyrus | L | 19 | -28 | -74 | 30 | 0.02 |  |
| Supp. Motor Area | R | 32 | 8 | 14 | 50 | 0.03 | 273 |
| Precentral Gyrus | L | 6 | -26 | -2 | 60 | 0.02 | 308 |
| Precentral Gyrus | L | 44 | -52 | 10 | 38 | 0.02 | 270 |
| Insular Cortex | L | 47 | -36 | 18 | -2 | 0.02 | 238 |
| Fusiform Gyrus | L | 37 | -42 | -54 | -18 | 0.02 | 166 |
| Inferior Parietal Lobe | R | 40 | 40 | -42 | 48 | 0.02 | 310 |
| Angular Gyrus | R | 7 | 32 | -66 | 48 | 0.02 |  |

**Note:** x, y, z: coordinates of the corresponding peak in MNI space, in units of millimeter.
Abbreviations: ALE, activation likelihood; BA, Brodmann area; K, cluster size in number of voxels; Regions are listed in a descending order by their peak Z value. For a cluster with multiple local peaks, the number of voxels in the whole cluster was only listed under the first local peak; L, left; R, right. <sup>a</sup> comprises the Frontal Eye Fields.

<u>The control</u> analysis of the 10 experiments reporting solely contrasts for the VMI *vs* Rest condition showed activation in precentral gyri and inferior parietal gyri, bilaterally, in the right superior parietal lobe and in the left supplementary motor area (**Figure 5** and **Table 6**). No activation emerged in early visual areas.

**Figure 5.**
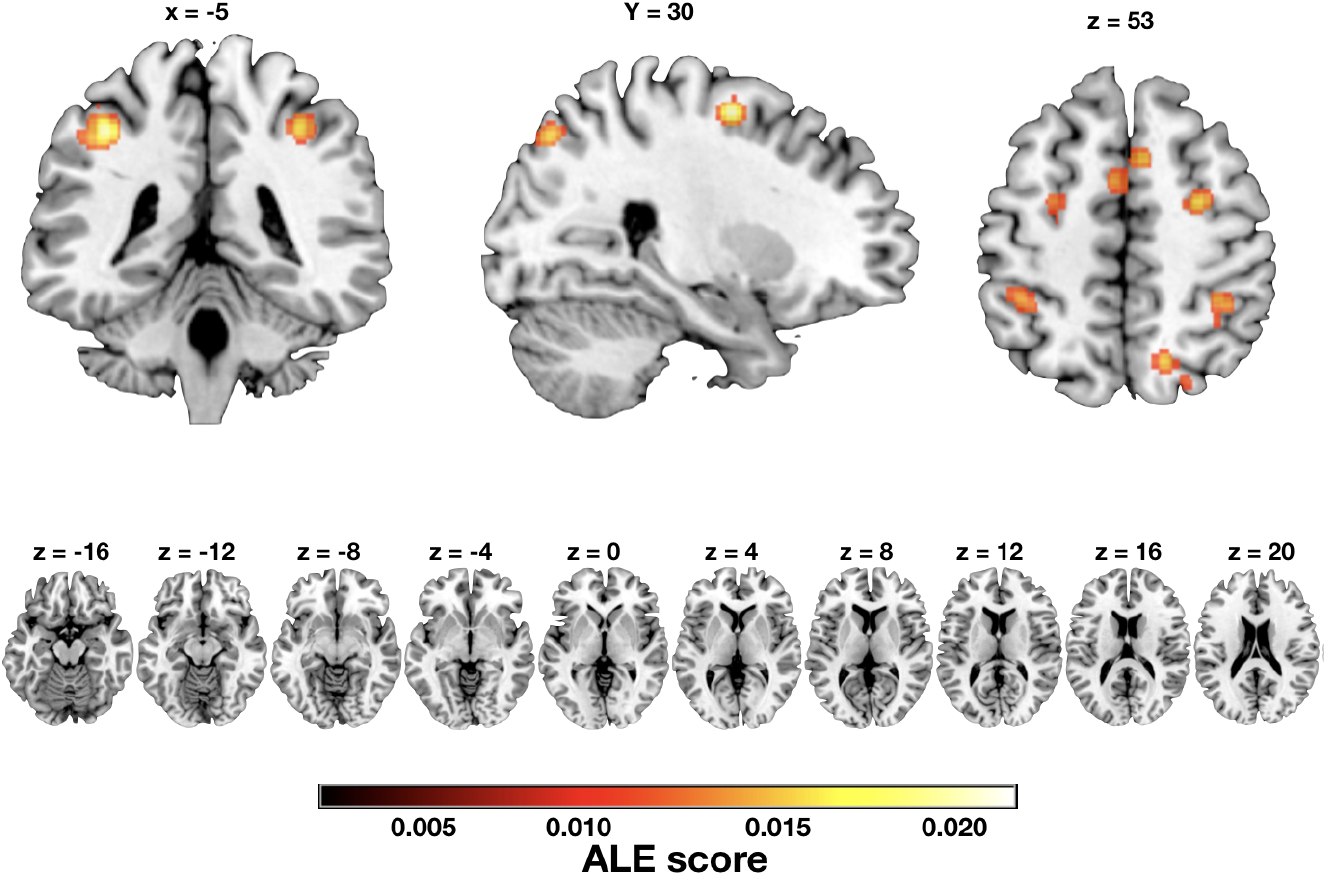
Brain regions showing consistently increased activation observed in the control analysis conducted on the VMI > Rest contrast, including fronto-parietal areas bilaterally.

**Table 6.** Brain regions showed consistent increased activation in the VMI *versus* Rest condition (10 studies, 131 foci, 144 subjects).

| Region | L/R | BA | x | y | z | Z | K |
| --- | --- | --- | --- | --- | --- | --- | --- |
| Precentral Gyrus | R | 6 | 28 | -4 | 56 | 0.02 | 146 |
| Inferior Parietal Lobe | L | 40 | -38 | -42 | 46 | 0.02 | 394 |
| Precentral Gyrus | L | 6 | -28 | -14 | 58 | 0.02 | 198 |
| Inferior Parietal Lobe | R | 40 | 36 | -42 | 48 | 0.02 | 170 |
| Superior Parietal Lobe | R | 7 | 16 | -66 | 52 | 0.01 | 143 |
| Suppl. Motor Area<sup>a</sup> | L | 6 | -4 | 10 | 52 | 0.01 | 88 |

**Note:** x, y, z: coordinates of the corresponding peak in MNI space, in units of millimeter.
Abbreviations: ALE, activation likelihood; BA, Brodmann area; K, cluster size in number of voxels; L, left; R, right. <sup>a</sup> This cluster extended bilaterally.

The meta-analysis of the 4 experiments included in the VMI *vs* Perception condition showed the activation increase in the insular cortex and in the supplementary motor area / anterior cingulate cortex, bilaterally. **Figure 6** shows the section views of the brain and highlights the regions with consistent activation increase in the VMI *vs* Perception condition, while **Table 7** reports the coordinates of common activation in the VMI *vs* Perception condition.

**Figure 6.**
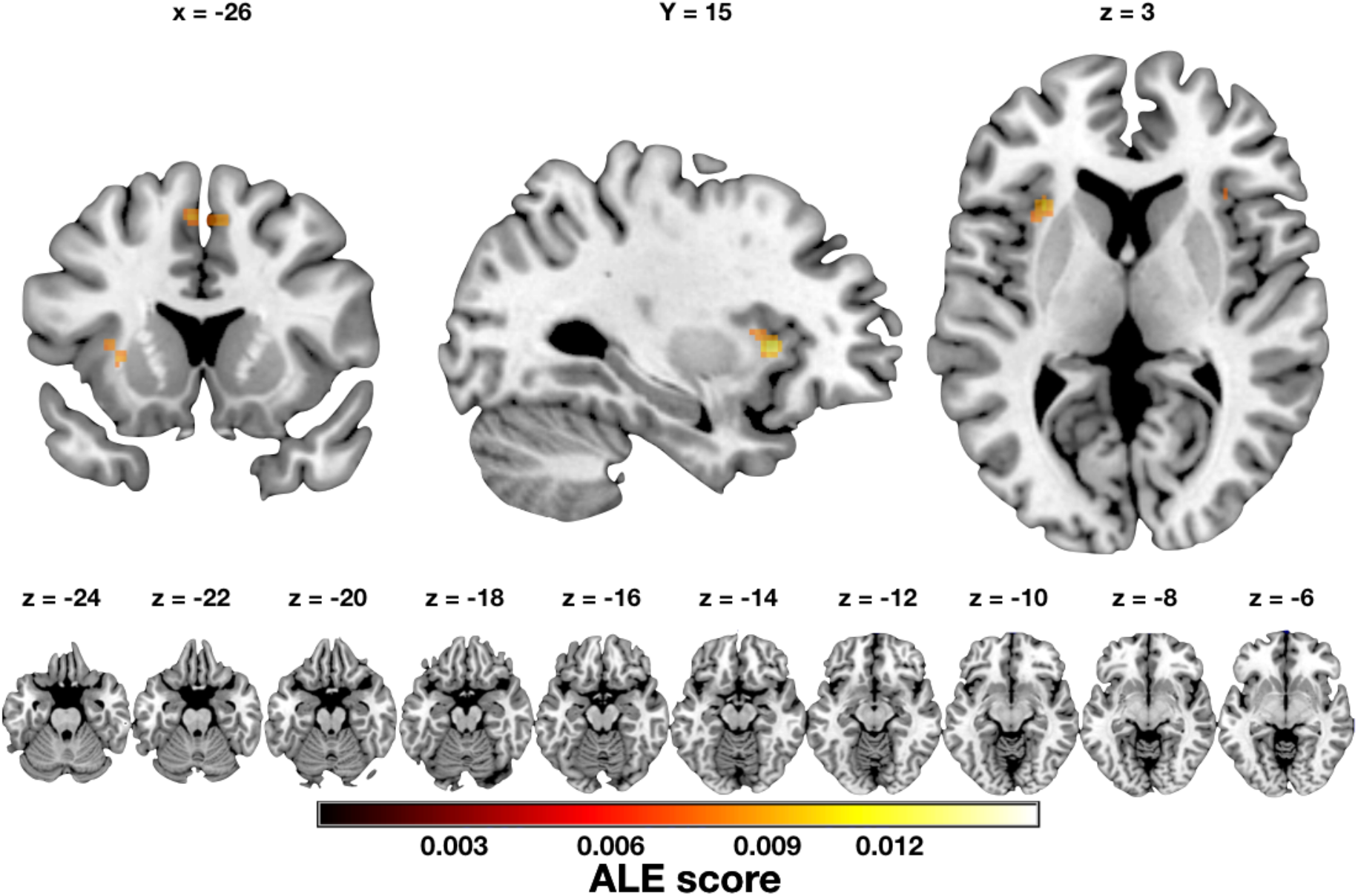
Brain regions showing consistent increased activation associated with the contrast VMI > Perception condition.

**Table 7.** Brain regions showed consistent increased activation in the VMI *versus* Perception condition (4 studies, 62 foci, 52 subjects).

| Region | L/R | BA | x | y | z | Z | K |
| --- | --- | --- | --- | --- | --- | --- | --- |
| Insular Cortex | R | 47 | 34 | 22 | 0 | 0.01 | 76 |
| Insular Cortex | L | 48 | -30 | 18 | 2 | 0.01 | 68 |
| Suppl. Motor Area<sup>a</sup> | L | 6 | -4 | 10 | 52 | 0.01 | 88 |

**Note:** x, y, z: coordinates of the corresponding peak in MNI space, in units of millimeter.
Abbreviations: ALE, activation likelihood; BA, Brodmann area; K, cluster size in number of voxels; L, left; R, right. <sup>a</sup> This cluster extended bilaterally.

The meta-analysis of the 15 experiments included in the MMI *versus* Control condition showed activation increases in the superior frontal gyri bilaterally, of the cerebellum bilaterally, and of the left supplementary motor area, of the anterior portion of the left calcarine sulcus, and of the parahippocampal / fusiform gyri. **Figure 7** shows the section views of the brain and highlights the regions with consistent activation increase MMI *vs* Control condition, while **Table 8** reports the coordinates of common activation in the MMI *vs* Control condition.

**Figure 7.**
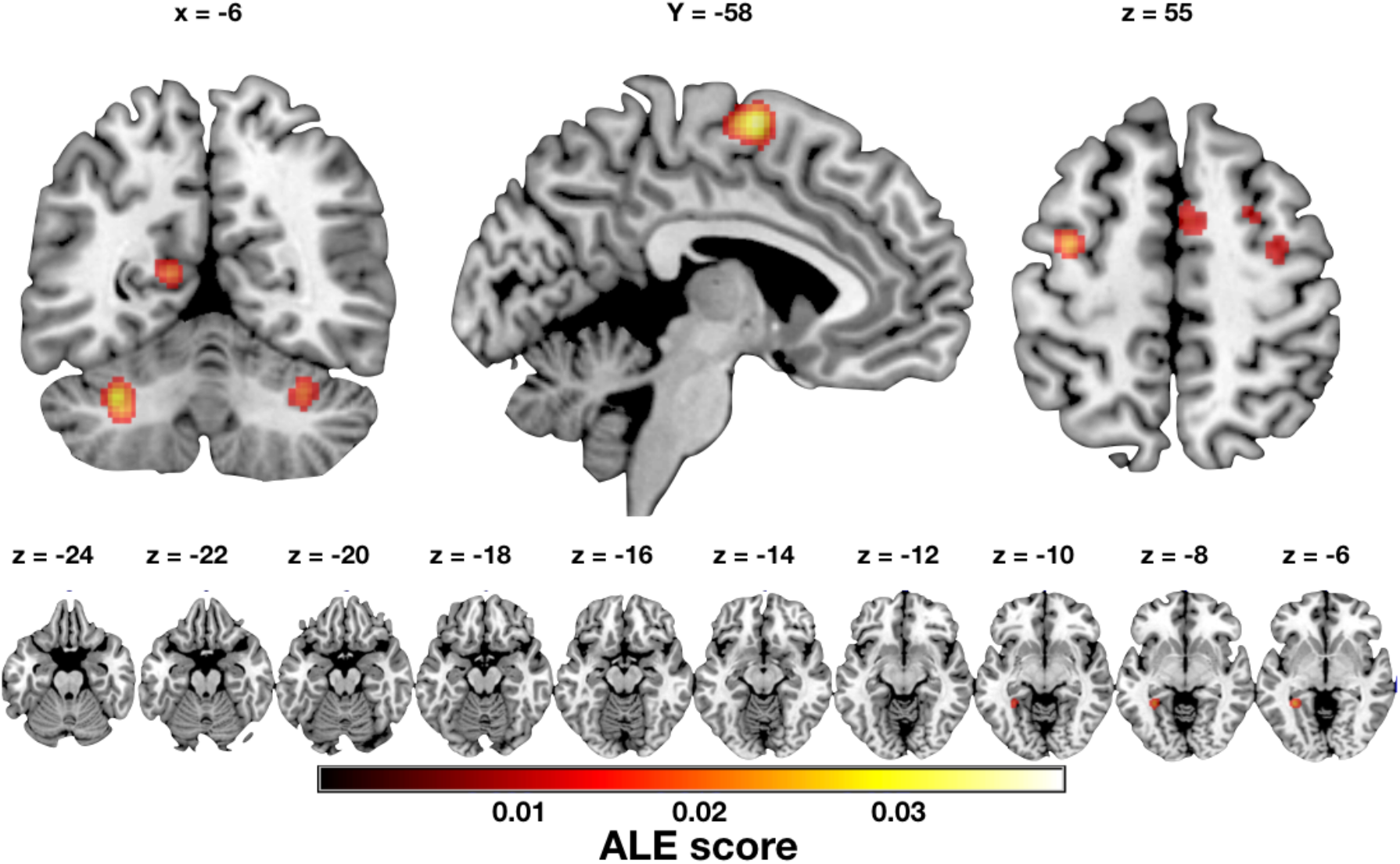
Brain regions showing consistent increased activation associated with the contrast MMI > Control condition.

**Table 8.** Brain regions showed consistent increased activation in the MMI *versus* Control condition contrast (15 studies, 340 foci, 239 subjects).

| Region | L/R | BA | x | y | z | Z | K |
| --- | --- | --- | --- | --- | --- | --- | --- |
| Supp. Motor Area | L | 6 | -2 | -2 | 66 | 0.04 | 604 |
| Cerebellum | L | 0 | -32 | -58 | -32 | 0.03 | 181 |
| Cerebellum | R | 0 | 32 | -62 | -30 | 0.03 | 135 |
| Precentral Gyrus <sup>a</sup> | L | 6 | -36 | -4 | 56 | 0.02 | 123 |
| Parahippocampal Gyrus | L | 37 | -32 | -44 | -4 | 0.02 | 100 |
| Calcarine Sulcus | L | 17 | -16 | -52 | 12 | 0.02 | 143 |
| Middle Frontal Gyrus <sup>a</sup> | R | 6 | 32 | 0 | 60 | 0.02 | 167 |

**Note:** x, y, z: coordinates of the corresponding peak in MNI space, in units of millimeter.
Abbreviations: ALE, activation likelihood; BA, Brodmann area; K, cluster size in number of voxels; L, left; R, right. <sup>a</sup> comprises the Frontal Eye Fields.

To further investigate common areas of activation among the VMIs contrasts, we conducted conjunction and disjunction analyses between the VMI *versus* Control and VMI *versus* Perception contrasts (31 experiments). Results of the conjunction analysis showed activation increases in the left anterior insular cortex and in the bilateral supplementary motor area. Results of both disjunction analyses showed no significant cluster of activation for the ((VMI > Control) > (VMI > Perception)) contrast. **Figure 8** shows the section views of the brain and highlights the regions with consistent activation increase in the conjunction analysis between the VMI > Control and VMI > Perception contrasts. No significant activation foci were found from either disjunction analysis. **Table 9** lists for the coordinates of common activation in the conjunction analyses.

**Figure 8.**
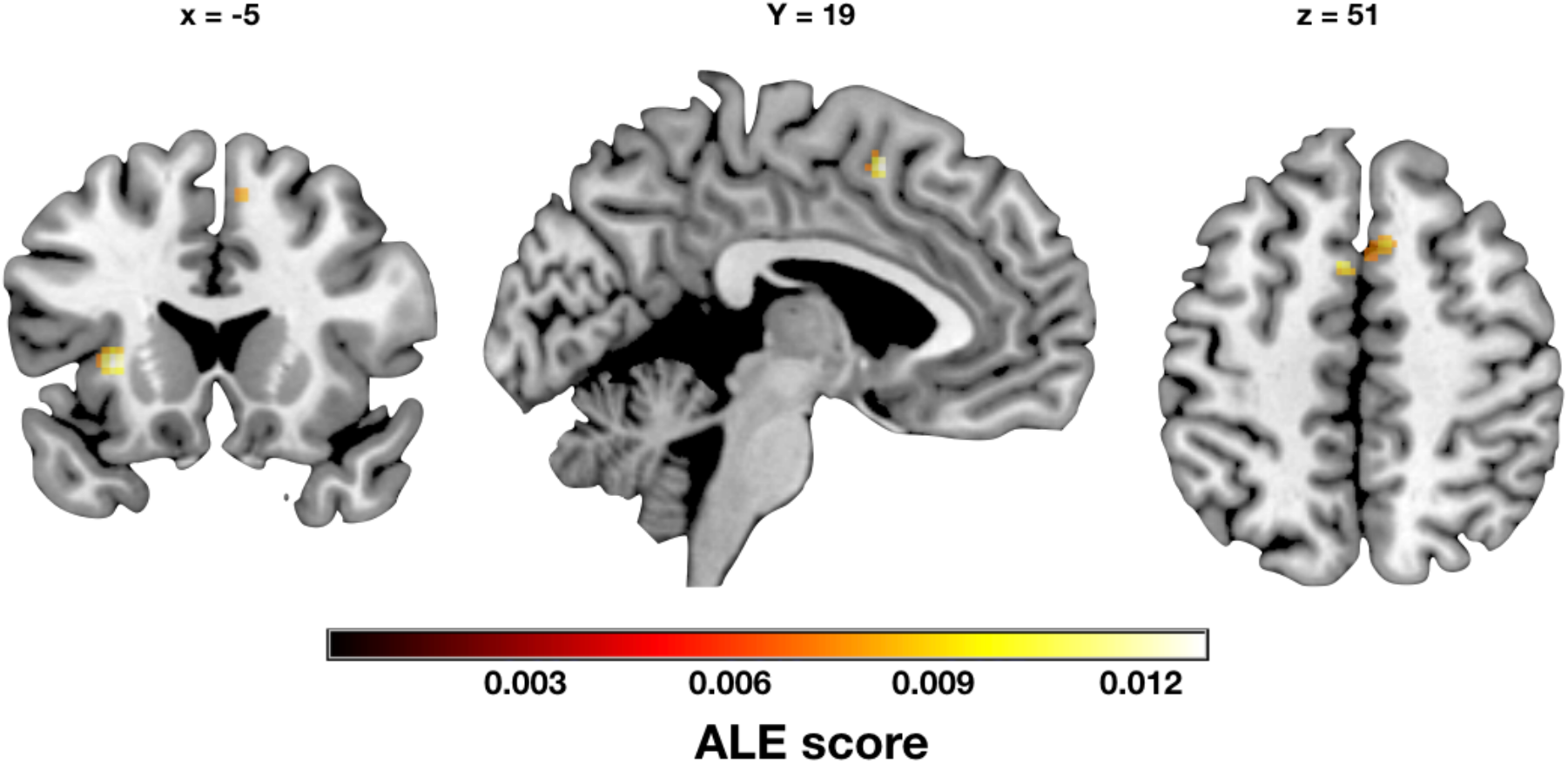
Brain regions showing consistent increased activation in the left anterior insular cortex and in the bilateral supplementary motor area / anterior cingulate cortex associated with the conjunction analysis between VMI versus Control condition and VMI versus Perception condition. See Table 9 for the coordinates of common activation in the conjunction analysis.

**Table 9.** Brain regions showed consistent increased activation in the conjunction and disjunction analyses between VMI *>* Control condition (27 studies, 376 foci, 380 subjects) and VMI > Perception condition (4 studies, 62 foci, 52 subjects).

| *Conjunction (VMI > Control) and (VMI > Perception)* |  |  |  |  |  |  |  |
| --- | --- | --- | --- | --- | --- | --- | --- |
| **Region** | **L/R** | **BA** | **x** | **y** | **z** | **Z** | **K** |
| Anterior Insular Cortex | L | 48 | -30 | 18 | 2 | 0.0156 | 248 |
| Supp. Motor Area<sup>a</sup> | L | 6 | -4 | 10 | 52 | 0.01 | 51 |
| *Disjunction (VMI > Control) > (VMI > Perception)* |  |  |  |  |  |  |  |
| **Region** | **L/R** | **BA** | **x** | **y** | **z** | **Z** | **K** |
| No cluster found |  |  |  |  |  |  |  |
| *Disjunction (VMI > Perception) > (VMI > Control)* |  |  |  |  |  |  |  |
| **Region** | **L/R** | **BA** | **x** | **y** | **z** | **Z** | **K** |
| No cluster found |  |  |  |  |  |  |  |

**Note:** x, y, z: coordinates of the corresponding peak in MNI space, in units of millimeter.
Abbreviations: ALE, activation likelihood; BA, Brodmann area; K, cluster size in number of voxels; L, left; R, right. <sup>a</sup> extending bilaterally.

The meta-analysis of the conjunction between the VMI *versus* Control and MMI *versus* Control contrasts (42 experiments) showed activation increases in the superior frontal gyri and in the supplementary motor area bilaterally. **Figure 9** shows the section views of the brain and highlights the regions with consistent activation increase in the conjunction and disjunction analyses between the VMI > Control and MMI > Control contrasts. See **Table 10** for the results of the conjunction and disjunction analyses.

**Figure 9.**
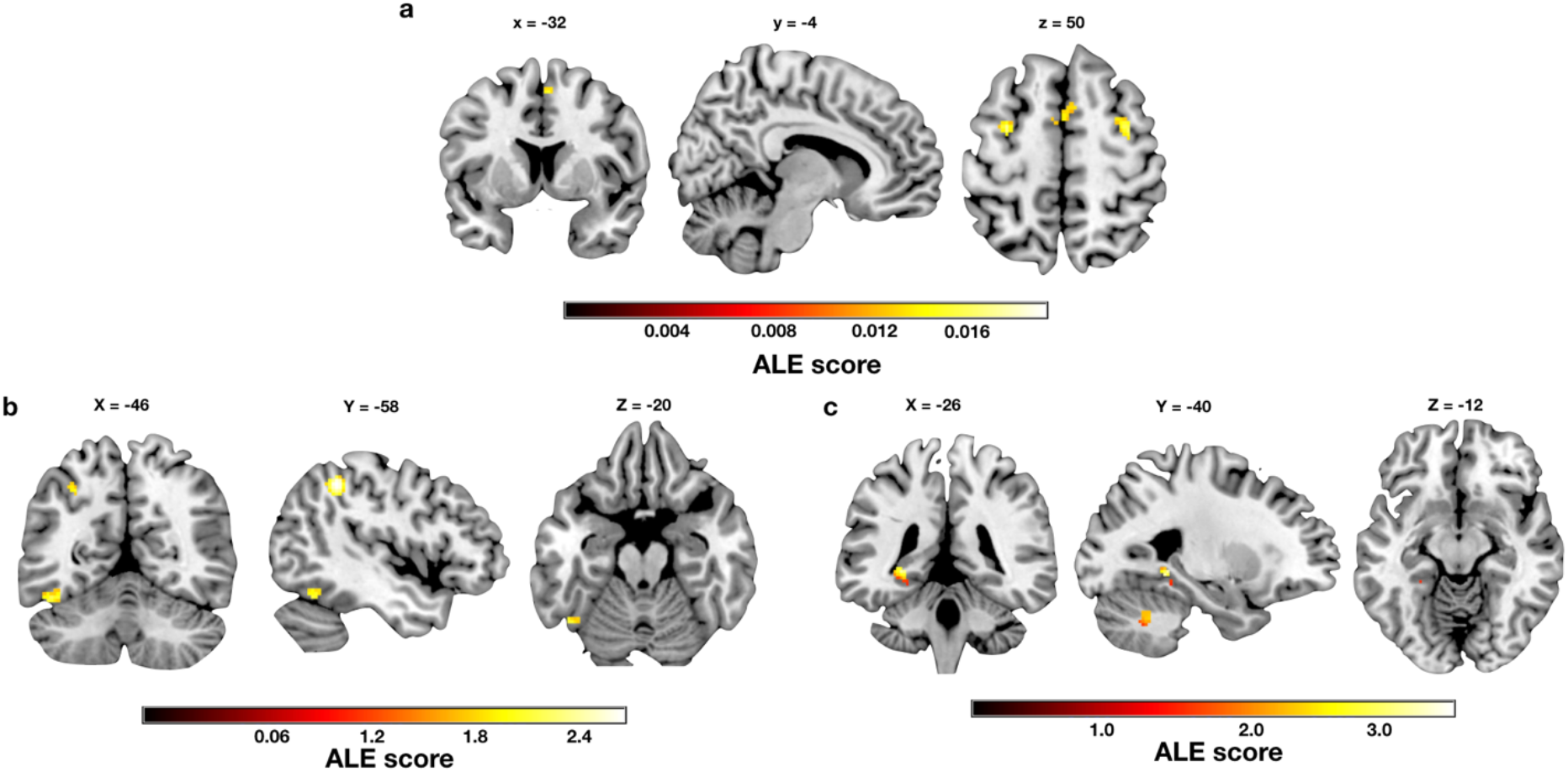
Brain regions showing consistent increased activation in the left anterior insular cortex and in the bilateral supplementary motor area / anterior cingulate cortex associated with the conjunction analysis between VMI versus Control condition and MMI versus Control condition. See Table 10 for the coordinates of common activation in the conjunction analysis.

**Table 10.** Brain regions showed consistent increased activation in the conjunction and disjunction analyses between VMI *>* Control condition (27 studies, 376 foci, 380 subjects) and MMI > Control condition (15 studies, 340 foci, 242 subjects).

| *Conjunction (VMI > Control) and (MMI > Control)* |  |  |  |  |  |  |  |
| --- | --- | --- | --- | --- | --- | --- | --- |
| **Region** | **L/R** | **BA** | **x** | **y** | **z** | **Z** | **K** |
| Precentral Gyrus | L | 6 | -32 | -4 | 50 | 3.00 | 54 |
| Precentral Gyrus | R | 6 | 34 | -8 | 52 | 2 | 56 |
| Supp. Motor Area <sup>a</sup> | L | 6 | -8 | 0 | 64 | 0.02 | 79 |
| *Disjunction (VMI > Control) > (MMI > Control)* |  |  |  |  |  |  |  |
| **Region** | **L/R** | **BA** | **x** | **y** | **z** | **Z** | **K** |
| Fusiform Gyrus | L | 37 | -46 | -58 | -20 | 3.00 | 60 |
| Superior Occipital Lobe | R | 7 | 28 | -78 | 44 | 2.00 | 61 |
| Inferior Parietal Lobe | L | 40 | -50 | -46 | 40 | 1.00 | 110 |
| *Disjunction (MMI > Control) > (VMI > Control)* |  |  |  |  |  |  |  |
| **Region** | **L/R** | **BA** | **x** | **y** | **z** | **Z** | **K** |
| Suppl. Motor Area | L | 6 | -10 | -8 | 70 | 13.00 | 450 |
| Suppl. Motor Area | R | 6 | 8 | 2 | 52 | 1.00 |  |
| Precentral Gyrus | L | 6 | -36 | -6 | 48 | 6.00 | 52 |
| Cerebellum Crus1 | R | 0 | 30 | -64 | -36 | 5.00 | 104 |
| Fusiform Gyrus | L | 37 | -26 | -40 | -12 | 4.00 | 106 |
| Cerebellum Crus1 | L | 0 | -32 | -58 | -38 | 3.00 | 151 |
| Lingual Gyrus | L | 18 | -14 | -56 | 4 | 2.00 | 157 |

**Note:** x, y, z: coordinates of the corresponding peak in MNI space, in units of millimeter.
Abbreviations: ALE, activation likelihood; BA, Brodmann area; K, cluster size in number of voxels; Regions are listed in a descending order by their peak Z value. For a cluster with multiple
local peaks, the number of voxels in the whole cluster was only listed under the first local peak;
L, left; R, right. <sup>a</sup> extending bilaterally.

**Figure 10** shows the substantial overlap between the VMI-related activation we found in the left fusiform gyrus and the cytoarchitectonic area FG4 (Lorenz et al., 2015), located in the rostral fusiform gyrus.

**Figure 10.**
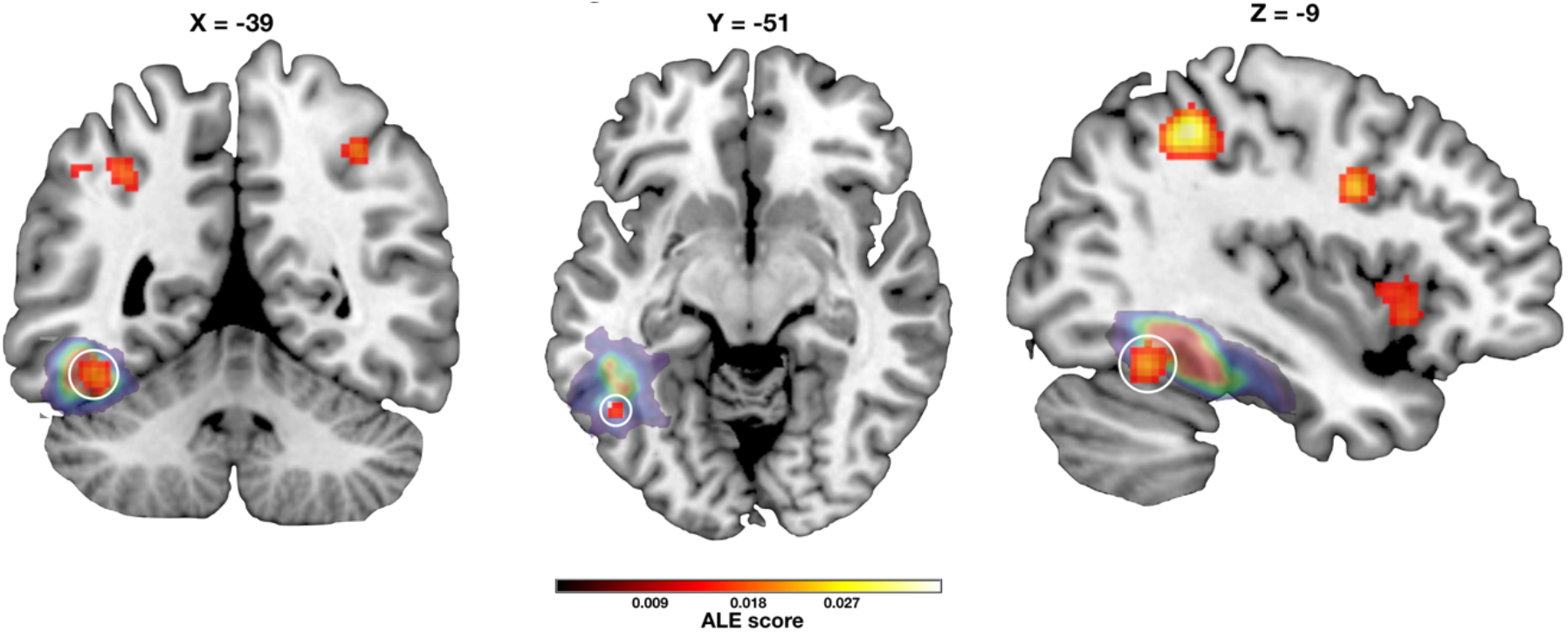
The left fusiform activation we found is colored in red and circled in white, while the mask of the probability distribution of the FG4 as shown in the coordinates reported by (Lorenz et al., 2015) is shown in colors ranging from purple to red.

### A Bayesian analysis of VMI studies

**Figure 11** shows the Bayes Factor (BF) values estimated using the spatial Bayesian latent factor regression model for the contrast VMI *vs* Control. Values in the early visual areas bilaterally were below the lower cut-off of 0.33, indicating the absence of consistent activation associated with VMI across the studies included in this meta-analysis.

**Figure 11.**
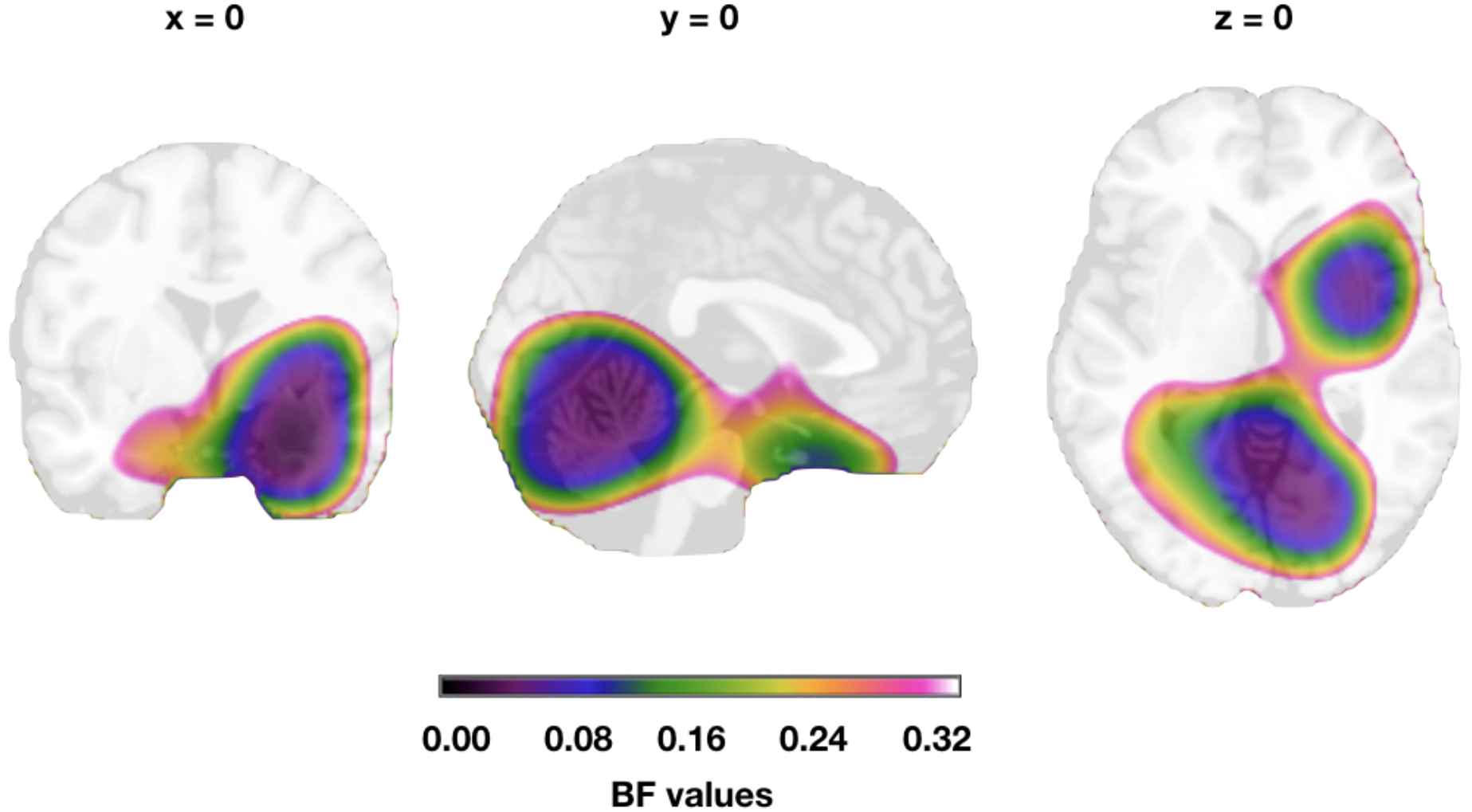
The area with color shading identifies BF values < 0.33, providing evidence for the absence of activation in that area. The area shaded in white corresponds to brain regions with BF values above 0.33.

## Discussion

### Lack of activation in primary sensory/motor cortices during mental imagery

The dominant model of VMI stipulates a functional and anatomical equivalence between VMI and visual perception (Pearson, 2019). According to this view, VMI and visual perception should rely upon the same cortical areas across the ventral cortical visual stream (Dijkstra et al., 2019; Pearson, 2019). Progression of visual information would occur in a bottom-up fashion during perception, and in a reversed, top-down direction during VMI. Indeed, results from hemodynamic neuroimaging in healthy volunteers demonstrate the engagement of early, occipital visual areas for VMI in some studies, but not in others. Notably, one study (Kilintari et al., 2016) actually showed a relative *decrease* of BOLD signal in occipital cortex during VMI. On the other hand, TMS interference on V1 was shown to impact VMI (Kosslyn et al., 1999), but this effect might have resulted from modulation of downstream visual areas. A TMS pulse applied over V1/V2 can stimulate not only local neuronal assemblies, but also excitatory projecting neurons reaching through the entire visual system, and beyond, up to FEF (Parkin et al., 2015; Bergmann and Hartwigsen, 2020). Also, more recent investigations showed mixed results. For example, Marzi et al. (2009) demonstrated different reaction times to TMS-induced and imagined phosphenes compared with those to phosphene-like dots presented on the screen. This is a critical difference between perception and imagery, that is specific to the effects of TMS on occipital areas. Keogh et al. (2020) reported that imagery strength showed a negative relationship with activation in early visual areas and a positive relationship with activation of frontal cortex clusters. Moreover, applying tDCS to decrease excitability in visual cortex, or to increase excitability in the superior frontal lobe, induced stronger mental images. Thus, neurostimulation evidence is at best ambiguous concerning the role of early visual areas in visual mental imagery, and suggests the involvement of superior frontal lobe areas, consistent with the FEF foci we found in the present meta-analysis.

The dominant model also provides a principled account of inter-individual differences in VMI, because the level of activation in early visual areas seem to correlate with the subjective experience of “vividness” of visual mental images (Cui et al., 2007). Recent support to this claim came from the finding that the content of VMI can be decoded from V1 activity (Senden et al., 2019). However, even the decoding studies are correlational and not causal in nature; their findings might thus reflect nonfunctional byproducts, instead of the true neural bases of VMI. In addition, a decoding study investigating face VMI (VanRullen and Reddy, 2019) reported that only voxels in the temporal lobe supported above-chance decoding of imagined faces, whereas occipital and frontoparietal regions did not perform above chance. Our Bayesian analysis indicates that we can confidently accept the hypothesis that VMI does not increase BOLD signal in occipital cortex when VMI is compared with control conditions. This result is unlikely to depend on different regions of V1 being implicated in different domains of VMI, because many of the included studies used items such as objects, words, and faces, which are preferentially processed in foveal vision. The present evidence, together with the extensive evidence from brain-damaged patients with intact VMI after lesions restricted to the occipital cortex (Bartolomeo, 2002), strongly suggests that VMI does not need activity in early visual areas. It might still be that some of the control tasks used in fMRI experiments might in fact have engaged visual mental imagery, contrary to the experimenters’ intention. This is a general methodological problem in the neuroimaging of visual mental imagery, and a similar argument might be made for the rest condition: the brain is never at rest, and we have no guarantee that subjects refrained from building mental images during “rest”, which typically induces mind wandering. Note that these possibilities would invalidate not only the present results, but virtually all the fMRI studies on visual mental imagery. To address this potential concern, we focused on the studies reporting an imagery>rest contrast (n= 10), although the power of this analysis may be limited. The results were consistent with the main analysis in confirming no visual cortex activation in the 10 studies included (Sack et al., 2002; Yomogida et al., 2004; Boly et al., 2007; Belardinelli et al., 2009; Soddu et al., 2009; Kaas et al., 2010; Seurinck et al., 2011; Sasaoka et al., 2014; Kilintari et al., 2016; Andersson et al., 2019) (see **Figure 5)**. In fact, only two of these studies reported an activation in early visual areas in the imagery > rest contrast (Boly et al., 2007; Kaas et al., 2010). We note, moreover, that at least some of the imagery > “rest and eyes closed” contrasts reported no activity whatsoever in V1 (Soddu et al., 2009 shows no occipital activation during rest and eyes closed; e.g., Kilintari et al., 2016 shows a decrease of activation in BA17 adn 18 during rest and eyes closed). More importantly, the absence of V1 activation demonstrated by our Bayesian analysis is highly consistent with abundant and converging evidence from studies on brain-damaged patients, who typically have intact visual mental imagery after lesions restricted to the occipital cortex. Most studies included here required participants to build mental images of complex objects, which were often retrieved from long-term memory. Future research should assess whether early visual areas may contribute to visual mental images of elementary forms (e.g., lines or gratings), or to mental images evoked from short-term memory (Ishai et al., 2000). Note, however, that some of the available evidence does not seem to support this possibility. Shelton and Pippitt (2006), found fMRI evidence of greater parietal and prefrontal activity in mental rotation compared to visual rotation. Broggin et al. (2012) found similar patterns of performance for luminance, contrast, and visual motion in visually presented and imagined stimuli, but important differences for frequency gratings.

We also note that for the motor imagery contrast we found no evidence of involvement of the primary motor cortex (BA4), consistent with previous meta-analytic results (Hétu et al., 2013). The lack of primary motor cortex activation might have been related to methodological differences in the tasks used in the studies included in the meta-analysis. Due to the topographical organization of BA4, different subregions, controlling different parts of the body, are meant to be activated by different tasks (e.g., drawing with the right hand should have activated the hand-portion of M1 in the left hemisphere, while a different region might have been activated for walking). Ultimately, this null effect may also reflect the obvious fact that actual movement execution has to be inhibited in motor mental imagery.

Results from the contrast between VMI and perception showed that the activation of the bilateral cingulo-opercular network (i.e., the anterior cingulate cortex and the anterior insular cortex) was greater in VMI than in perception. This result should, however, be interpreted with caution, given the low number of studies that fit our inclusion and exclusion criteria for the VMI > Perception contrast (n = 4).

### Left fusiform involvement in VMI

The present demonstration of robust activity in the left fusiform gyrus during VMI is remarkable in view of its substantial overlapping with the cytoarchitectonic area FG4 (see Fig. 2), despite the wide variety of methods used in the included studies, and because of its striking agreement with the evidence from brain-damaged patients. Impaired VMI in these patients typically occurs after extensive damage to the temporal lobe, and especially in the left hemisphere, at least for form and color VMI (Bartolomeo, 2002, 2008; Moro et al., 2008). Although anatomical evidence is rarely precise in stroke patients, a recently described case report (Thorudottir et al., 2020) provides additional, converging causal evidence on the role of the left fusiform gyrus in VMI. After a bilateral stroke in the territory of the posterior cerebral artery, an architect, who before the stroke could easily imagine objects and buildings, spontaneously reported to have become virtually unable to visualize items. He had now to rely on computer-aided design for his work, because he needed to see items that he could previously imagine. The stroke had provoked extensive damage to the right hemisphere, including the occipital pole, the lingual gyrus, the fusiform gyrus and the parahippocampal region. In the left hemisphere, the lesion was smaller and affected only the medial fusiform gyrus and lingual gyrus. Comparison of the lesion location with those of other patients with strokes in the same arterial territory, but spared VMI, showed that the patient with impaired VMI had selective damage in the right lingual gyrus and in the left posterior medial fusiform gyrus. The fusiform lesion was located in close proximity to the hotspot we found in the present meta-analysis. This fusiform region might act as an interface between semantic processing in the anterior temporal lobe (Lambon Ralph et al., 2017) and perceptual information coming from the occipital cortex. There is, however, the possibility that VMI-related regions in the left temporal lobe are more extended than suggested by the present fMRI results. BOLD responses in the anterior temporal lobe suffer from large magnetic susceptibility artifacts arising from the auditory canals (Wandell, 2011). Even when distortion-corrected sequences are used in fMRI, the lowest signal is recorded from the region between the midfusiform region and the temporal pole. Thus, additional, more anterior temporal regions could potentially participate in VMI-related abilities.

### Fronto-parietal networks and VMI

The bilateral activation of areas within the fronto-parietal networks (Corbetta, 1998; Rossi et al., 2009; Xuan et al., 2016) and the cingulo-opercular network (Dosenbach et al., 2008; Sadaghiani and D’Esposito, 2015; Sheffield et al., 2015; Dubis et al., 2016) suggests that mental imagery requires the activation of task-positive neural substrates supporting high-level cognitive functions, including those for attentional control (Petersen and Posner, 2012; Xuan et al., 2016; Bartolomeo and Seidel Malkinson, 2019; Wu et al., 2019) and visual working memory (Awh and Jonides, 2001; Jonikaitis and Moore, 2019). Also, the implication of fronto-parietal networks in VMI is broadly consistent with lesion location in spatial neglect for visual mental images (Bisiach and Luzzatti, 1978), which typically goes far beyond the occipital cortex, and implicates the fronto-parietal networks in the right hemisphere (Guariglia et al., 1993; Bartolomeo et al., 1994; Rode et al., 2010). Taken together, the results from this meta-analysis shed further light on the neural correlates of VMI and suggest a left-hemisphere superiority that is in line with neuropsychological evidence (Basso et al., 1980; Riddoch, 1990; Goldenberg, 1992; Moro et al., 2008). Functional lateralization is a fundamental organization principle of the brain, and spans across the anatomical and functional realm (Karolis et al., 2019). Besides adding additional evidence regarding the potential lateralization of VMI, this result can be put in relation to attention processes, another family of functions showing various types of brain lateralization (Heilman and Van Den Abell, 1980; Bartolomeo, 2007; Asanowicz et al., 2012; Chica et al., 2012; Bartolomeo, 2014; Spagna et al., 2016; Spagna et al., 2018; Bartolomeo and Seidel Malkinson, 2019). Putting together the high degree of similarity between the regions of the brain showing a consistent increase of activation associated with VMI and those associated with visual attention, it is likely that these two functions share a common bilateral neural substrate in the rostral frontal regions; a right hemisphere bias might emerge when directing attention towards external stimuli (as in visual cued tasks) (Bartolomeo and Seidel Malkinson, 2019), while a left hemisphere bias might arise when directing attention towards internally-generated stimuli (Gazzaley and Nobre, 2012) (as in the case of mental imagery).

Given the prominent role of fronto-parietal attention networks in attention-mediated conscious processing (Chica and Bartolomeo, 2012; Chica et al., 2013; Chica et al., 2014) and in neural models of consciousness (Dehaene and Changeux, 2011), it is tempting to speculate that individual differences in subjective vividness of visual mental images might in part depend on different degrees of engagement of fronto-parietal networks in VMI, or on their degree of anatomical and functional connectivity with the ventral temporal cortex. For example, vivid reexperiencing of traumatic memories in post-traumatic stress disorder might depend in part on poor prefrontal control on medial temporal lobe activity (Mary et al., 2020).

### A revised neural model of VMI

The present results, together with the evidence from neurological patients (Bartolomeo et al., 2020), invite a revision of the neural model of VMI (**Figure 12**).

**Figure 12.**
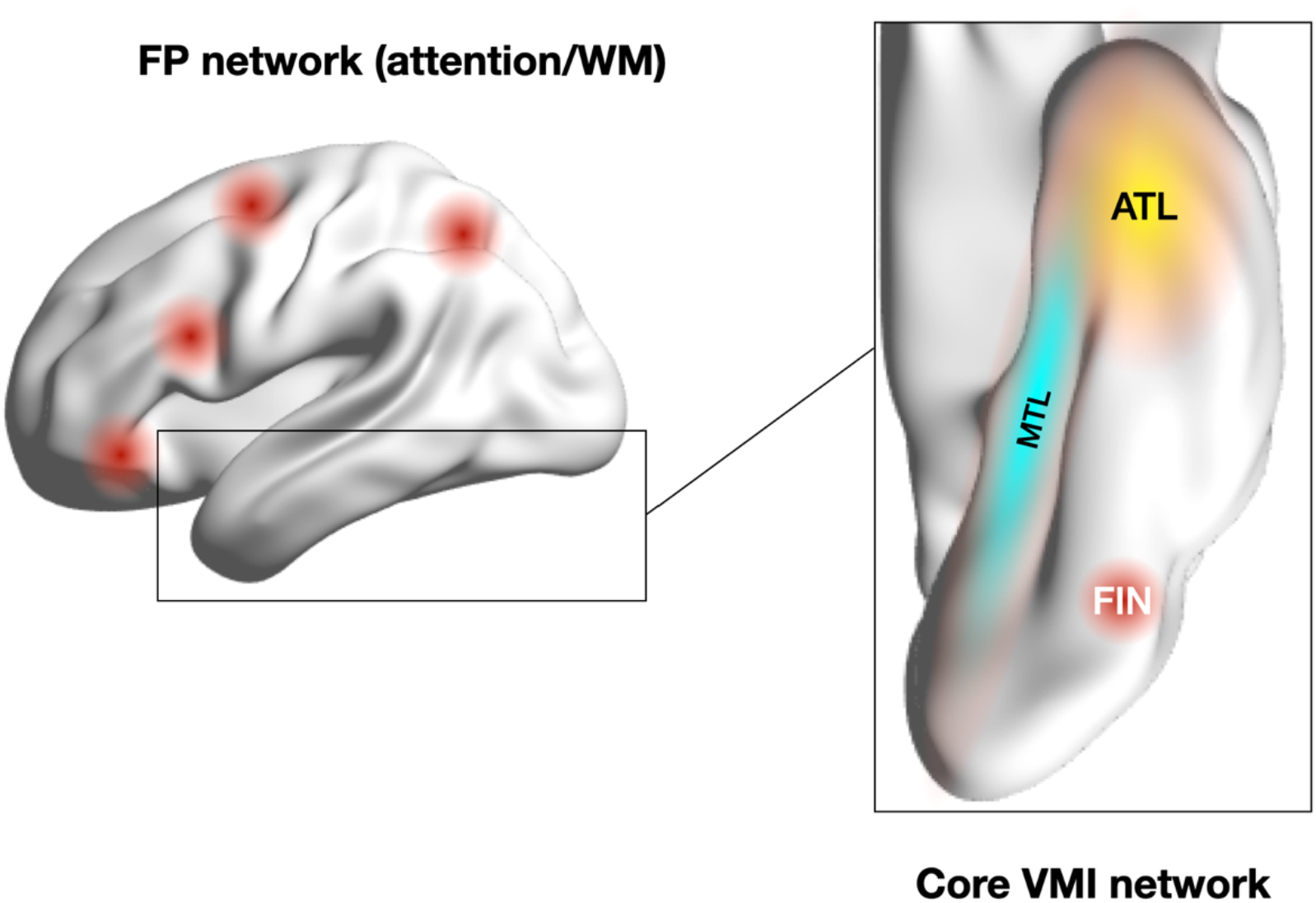
Schematic depiction of a revised neural model of VMI. FP, fronto-parietal; WM, visual Working Memory; FIN, Fusiform Imagery Node; ATL, Anterior Temporal Lobe; MTL; Medial Temporal Lobe.

We propose that a core network of VMI builds upon a region in the FG4 area (Lorenz et al., 2015) of the left fusiform gyrus, which may be labeled as Fusiform Imagery Node (FIN), in analogy with other domain-preferring regions in the ventral temporal cortex (Mahon and Caramazza, 2011). This region corresponds roughly to the lateral section of left middle fusiform region, in the vicinity of regions preferentially responding to letter strings (Cohen et al., 2002) and to faces (although predominantly in the right hemisphere (Kanwisher et al., 1997) (but also see Bukowski et al., 2013 for a focus on the left fusiform gyrus). More anterior regions of the left temporal lobe (ATL) provide the FIN with semantic information (Lambon Ralph et al., 2017). Importantly, both left and right hemisphere ATL are implicated in conceptual knowledge (Lambon Ralph et al., 2017). This bilateral organization might allow some patients with unilateral brain damage to compensate for deficits of VMI (see Bartolomeo and de Schotten, 2016) As noted above, we cannot exclude the participation of more anterior regions in the ventral temporal cortex, between the mid-fusiform gyrus and the temporal pole, which are difficult to visualize in fMRI because of susceptibility artifacts. During VMI, prefrontal circuits might engage activity in this core ventral temporal network. The FIN is part of the ventral cortical visual pathway; during perception, it receives visual information from more posterior regions. Medial temporal lobe (MTL) structures may provide a neural substrate for recombining elements of past experiences to generate episodic simulations in VMI (Schacter et al., 2007; Mahr, 2020). MTL structures, together with the posterior cingulate, and perhaps with the fronto-parietal attention networks, may contribute to the phenomenal experience of VMI as “quasi-visual” in nature (sometimes referred to as “vividness” of visual mental images) (Fulford et al., 2018). The degree of this integration may differ among individuals. Functional disconnection of the MTL components of the network may lead to impaired phenomenal experience associated with VMI (aphantasia). In these cases, some recovery of visual information (without VMI experience) may still be provided by the ATL-FIN portion of the VMI network. Dysfunction of the ATL-FIN component may lead to VMI deficits, as suggested by the patterns of performance of patients with extensive left temporal damage (Bartolomeo, 2002). Domain-specific VMI might show differences in hemispheric laterality. For example, a homologue network in the right hemisphere might contribute to aspects of VMI for faces (O’Craven and Kanwisher, 2000; Barton and Cherkasova, 2003; Sunday et al., 2018) (but also see Ishai et al., 2000), in analogy with the previously mentioned right fusiform superiority in perceptual face processing (Kanwisher et al., 1997; but also see Bukowski et al., 2013). The present evidence is consistent with the possibility that prefrontal regions initiate visual mental imagery, in analogy with other endogenous cognitive processes, such as top-down spatial attention (Buschman and Miller, 2007). In our model, VMI-related activity in the temporal lobe core network is initiated, modulated and main tained by fronto-parietal networks subserving attention and working memory. Dysfunction of right hemisphere fronto-parietal networks, coupled with their disconnection from the left FIN (Rode et al., 2010), but with normal functioning of the left hemisphere portions of the network, may provoke neglect for the left side part of visual mental images, with spared processing of their right side (imaginal or representational neglect) (Bisiach and Luzzatti, 1978; Guariglia et al., 1993; Bartolomeo et al., 1994). In this case, visual mental images are built by the left hemisphere VMI network, but the attentional impairment resulting from right fronto-parietal dysfunction determines the incomplete exploration or maintenance of the left portion of mental images (Rode et al., 2010).

## Conclusions

Altogether, the long-standing neuropsychological evidence of the past 30 years, together with more recent neuroimaging evidence analyzed here, invites a reappraisal of the dominant model of visual mental imagery, which was based on a substantial functional and structural overlap between visual mental imagery and visual perception. The available evidence suggests a different scenario, where prefrontal regions initiate the imagination process by recruiting the stored information to-be-recalled from semantic circuits involving the left temporal lobe, together with high-level ventral stream areas of the left fusiform and parahippocampal gyri, and then activates networks important for attention and visual working memory, which maintain the recalled images to perform the task at hand (whether a simple inspection, a rotation, or a judgement). This alternative model seems better equipped to deal with dramatic patterns of double dissociation between perception and imagery abilities in brain-damaged patients. In particular, the present finding of FIN activity in VMI is strikingly consistent with the observation of imagery deficits occurring not after occipital damage, but after extensive left temporal damage (Bartolomeo, 2002). Further neuroimaging and neuropsychological research is needed to determine the brain correlates of specific domains of VMI (e.g., faces, object form or color, places, orthographic material). A clear description of these fast-occurring dynamics can only be achieved by employing techniques with adequate spatiotemporal resolution, such as intracerebral recordings (see e.g., Rossion et al., 2018), to trace feedback and feedforward sweeps of activation in the present model.

## Authors Contribution

P.B. and A.S. conceived the study. A.S., D.H., and J.L. extracted the coordinates from the source articles. A.S. and D.H. conducted the analysis on Ginger ALE. A.S. and J.L. conducted the Bayesian Analyses. All the authors contributed to the writing of the manuscript. Raw Coordinates included in each of the contrasts, the maps of the results in the .nii format as well as the raw outputs from GingerALE software, code and instructions to reproduce the Bayes analyses, instructions to reproduce the figures, and supplementary videos can be found on the Github page of A.S.

## Acknowledgments

Supported by an ICM post-doctoral fellowship to A.S., and by funding from Dassault Systèmes. P.B. is supported by the Agence Nationale de la Rercherche through ANR-16-CE37-0005 and ANR-10-IAIHU-06. We thank Laurent Cohen, Tal Seidel Malkinson, Silvia Montagna, Katarzyna Siuda-Krzywicka, Michel Thiebaut de Schotten, and an anonymous reviewer for helpful discussion.

## Reference List

Aglioti S, Bricolo E, Cantagallo A, Berlucchi G (1999) Unconscious letter discrimination is enhanced by association with conscious color perception in visual form agnosia. Current Biology 9:1419–1422.

Andersson P, Ragni F, Lingnau A (2019) Visual imagery during real-time fMRI neurofeedback from occipital and superior parietal cortex. NeuroImage 200:332–343.

Asanowicz D, Marzecova A, Jaskowski P, Wolski P (2012) Hemispheric asymmetry in the efficiency of attentional networks. Brain Cogn 79:117–128.

Awh E, Jonides J (2001) Overlapping mechanisms of attention and spatial working memory. Trends in cognitive sciences 5:119–126.

Bartolomeo P (2002) The relationship between visual perception and visual mental imagery: a reappraisal of the neuropsychological evidence. Cortex 38:357–378.

Bartolomeo P (2007) Visual neglect. Current opinion in neurology 20:381–386.

Bartolomeo P (2008) The neural correlates of visual mental imagery: An ongoing debate. Cortex 44:107–108.

Bartolomeo P (2014) The attention systems of the human brain. In: Attention Disorders After Right Brain Damage, pp 1–19: Springer London.

Bartolomeo P, de Schotten MT (2016) Let thy left brain know what thy right brain doeth: Interhemispheric compensation of functional deficits after brain damage. Neuropsychologia 93:407–412.

Bartolomeo P, Seidel Malkinson T (2019) Hemispheric lateralization of attention processes in the human brain. Current Opinion in Psychology 29C:90–96.

Bartolomeo P, D’Erme P, Gainotti G (1994) The relationship between visuospatial and representational neglect. Neurology 44:1710–1714.

Bartolomeo P, Bourgeois A, Bourlon C, Migliaccio R (2013) Visual and motor mental imagery after brain damage. In: Multisensory Imagery (Lacey S, Lawson R, eds), pp 249–269. New York: Springer.

Bartolomeo P, Hajhajate D, Liu J, Spagna A (2020) Assessing the causal role of early visual areas in visual mental imagery. Nature Reviews Neuroscience 21:517–517.

Barton JJ, Cherkasova M (2003) Face imagery and its relation to perception and covert recognition in prosopagnosia. Neurology 61:220–225.

Basso A, Bisiach E, Luzzatti C (1980) Loss of mental imagery: A case study. Neuropsychologia 18:435–442.

Behrmann M, Winocur G, Moscovitch M (1992) Dissociation between mental imagery and object recognition in a brain-damaged patient. Nature 359:636.

Belardinelli MO, Palmiero M, Sestieri C, Nardo D, Di Matteo R, Londei A, D’Ausilio A, Ferretti A, Del Gratta C, Romani GL (2009) An fMRI investigation on image generation in different sensory modalities: the influence of vividness. Acta psychologica 132:190–200.

Bergmann TO, Hartwigsen G (2020) Inferring Causality from Noninvasive Brain Stimulation in Cognitive Neuroscience. Journal of Cognitive Neuroscience:1–29.

Beschin N, Cocchini G, Della Sala S, Logie RH (1997) What the eyes perceive, the brain ignores: a case of pure unilateral representational neglect. Cortex 33:3–26.

Bisiach E, Luzzatti C (1978) Unilateral neglect of representational space. Cortex 14:129–133.

Bisiach E, Luzzatti C, Perani D (1979) Unilateral neglect, representational schema and consciousness. Brain: a journal of neurology 102:609–618.

Boly M, Coleman MR, Davis M, Hampshire A, Bor D, Moonen G, Maquet PA, Pickard JD, Laureys S, Owen AM (2007) When thoughts become action: an fMRI paradigm to study volitional brain activity in non-communicative brain injured patients. NeuroImage 36:979–992.

Broggin E, Savazzi S, Marzi CA (2012) Similar effects of visual perception and imagery on simple reaction time. Quarterly Journal of Experimental Psychology 65:151–164.

Bukowski H, Dricot L, Hanseeuw B, Rossion B (2013) Cerebral lateralization of face-sensitive areas in left-handers: only the FFA does not get it right. Cortex 49:2583–2589.

Buschman TJ, Miller EK (2007) Top-down versus bottom-up control of attention in the prefrontal and posterior parietal cortices. science 315:1860–1862.

Chatterjee A, Southwood MH (1995) Cortical blindness and visual imagery. Neurology 45:2189–2195.

Chica AB, Bartolomeo P (2012) Attentional routes to conscious perception. Frontiers in Psychology 3:1–12.

Chica AB, Paz-Alonso PM, Valero-Cabre A, Bartolomeo P (2013) Neural bases of the interactions between spatial attention and conscious perception. Cerebral Cortex 23:1269–1279.

Chica AB, Valero-Cabre A, Paz-Alonso PM, Bartolomeo P (2014) Causal contributions of the left frontal eye field to conscious perception. Cerebral Cortex 24:745–753.

Chica AB, Thiebaut de Schotten M, Toba M, Malhotra P, Lupianez J, Bartolomeo P (2012) Attention networks and their interactions after right-hemisphere damage. Cortex 48:654–663.

Cohen L, Lehéricy S, Chochon F, Lemer C, Rivaud S, Dehaene S (2002) Language-specific tuning of visual cortex? Functional properties of the Visual Word Form Area. Brain: a journal of neurology 125:1054–1069.

Corbetta M (1998) Frontoparietal cortical networks for directing attention and the eye to visual locations: identical, independent, or overlapping neural systems? Proceedings of the National Academy of Sciences of the United States of America 95:831–838.

Cui X, Jeter CB, Yang D, Montague PR, Eagleman DM (2007) Vividness of mental imagery: individual variability can be measured objectively. Vision research 47:474–478.

de Gelder B, Tamietto M, Pegna AJ, Van den Stock J (2015) Visual imagery influences brain responses to visual stimulation in bilateral cortical blindness. Cortex 72:15–26.

de Vito S, Bartolomeo P (2016) Refusing to imagine? On the possibility of psychogenic aphantasia. A commentary on Zeman et al. (2015). Cortex 74:334–335.

Dehaene S, Changeux JP (2011) Experimental and theoretical approaches to conscious processing. Neuron 70:200–227.

Dentico D, Cheung BL, Chang JY, Guokas J, Boly M, Tononi G, Van Veen B (2014) Reversal of cortical information flow during visual imagery as compared to visual perception. NeuroImage 100:237–243.

Dijkstra N, Bosch SE, van Gerven MA (2017a) Vividness of Visual Imagery Depends on the Neural Overlap with Perception in Visual Areas. The Journal of Neuroscience 37:1367–1373.

Dijkstra N, Bosch SE, van Gerven MAJ (2019) Shared Neural Mechanisms of Visual Perception and Imagery. Trends in cognitive sciences 23:423–434.

Dijkstra N, Zeidman P, Ondobaka S, van Gerven MAJ, Friston K (2017b) Distinct Top-down and Bottom-up Brain Connectivity During Visual Perception and Imagery. Scientific reports 7:5677.

Dijkstra N, Mostert P, Lange FP, Bosch S, van Gerven MA (2018) Differential temporal dynamics during visual imagery and perception. eLife 7.

Dosenbach, Fair DA, Cohen AL, Schlaggar BL, Petersen SE (2008) A dual-networks architecture of top-down control. Trends in cognitive sciences 12:99–105.

Dubis JW, Siegel JS, Neta M, Visscher KM, Petersen SE (2016) Tasks driven by perceptual information do not recruit sustained BOLD activity in cingulo-opercular regions. Cerebral Cortex 26:192–201.

Eickhoff SB, Bzdok D, Laird AR, Kurth F, Fox PT (2012) Activation likelihood estimation metaanalysis revisited. NeuroImage 59:2349–2361.

Eickhoff SB, Laird AR, Grefkes C, Wang LE, Zilles K, Fox PT (2009) Coordinate-based activation likelihood estimation meta-analysis of neuroimaging data: A random-effects approach based on empirical estimates of spatial uncertainty. Human brain mapping 30:2907–2926.

Farah MJ, Levine DN, Calvanio R (1988) A case study of mental imagery deficit. Brain and cognition 8:147–164.

Fulford J, Milton F, Salas D, Smith A, Simler A, Winlove C, Zeman A (2018) The neural correlates of visual imagery vividness–An fMRI study and literature review. Cortex 105:26–40.

Galton F (1880) Visualised numerals. Nature 21:252–256.

Gazzaley A, Nobre AC (2012) Top-down modulation: bridging selective attention and working memory. Trends in cognitive sciences 16:129–135.

Goldenberg G (1992) Loss of visual imagery and loss of visual knowledge—a case study. Neuropsychologia 30:1081–1099.

Goldenberg G, Mullbacher W, Nowak A (1995) Imagery without perception – A case study of anosognosia for cortical blindness. Neuropsychologia 33:1373–1382.

Guariglia C, Padovani A, Pantano P, Pizzamiglio L (1993) Unilateral neglect restricted to visual imagery. Nature 364:235–237.

Heilman KM, Van Den Abell T (1980) Right hemisphere dominance for attention The mechanism underlying hemispheric asymmetries of inattention (neglect). Neurology 30:327–327.

Hétu S, Grégoire M, Saimpont A, Coll M-P, Eugène F, Michon P-E, Jackson PL (2013) The neural network of motor imagery: An ALE meta-analysis. Neuroscience & Biobehavioral Reviews 37:930–949.

Ibáñez-Marcelo E, Campioni L, Phinyomark A, Petri G, Santarcangelo EL (2019) Topology highlights mesoscopic functional equivalence between imagery and perception: The case of hypnotizability. NeuroImage 200:437–449.

Ishai A, Ungerleider LG, Haxby JV (2000) Distributed neural systems for the generation of visual images. Neuron 28:979–990.

Jacobs C, Schwarzkopf DS, Silvanto J (2018) Visual working memory performance in aphantasia. Cortex 105:61–73.

Jonikaitis D, Moore T (2019) The interdependence of attention, working memory and gaze control: behavior and neural circuitry. Current Opinion in Psychology 29:126–134.

Kaas A, Weigelt S, Roebroeck A, Kohler A, Muckli L (2010) Imagery of a moving object: the role of occipital cortex and human MT/V5+. NeuroImage 49:794–804.

Kanwisher N, McDermott J, Chun MM (1997) The fusiform face area: a module in human extrastriate cortex specialized for face perception. Journal of neuroscience 17:4302–4311.

Karolis VR, Corbetta M, Thiebaut De Schotten M (2019) The architecture of functional lateralisation and its relationship to callosal connectivity in the human brain. Nature communications 10:1417.

Keogh R, Bergmann J, Pearson J (2020) Cortical excitability controls the strength of mental imagery. eLife 9:e50232.

Kilintari M, Narayana S, Babajani-Feremi A, Rezaie R, Papanicolaou AC (2016) Brain activation profiles during kinesthetic and visual imagery: An fMRI study. Brain Research 1646:249–261.

Kosslyn SM, Ganis G, Thompson WL (2001) Neural foundations of imagery. Nat Rev Neurosci 2:635–642.

Kosslyn SM, Thompson WL, Ganis G (2006) The case for mental imagery: Oxford University Press.

Kosslyn SM, Pascual-Leone A, Felician O, Camposano S, Keenan JP, Thompson WL, Ganis G, Sukel KE, Alpert NM (1999) The role of area 17 in visual imagery: convergent evidence from PET and rTMS. Science 284:167–170.

Lambon Ralph MA, Jefferies E, Patterson K, Rogers TT (2017) The neural and computational bases of semantic cognition. Nature Reviews Neuroscience 18:42.

Lee S-H, Kravitz DJ, Baker CI (2012) Disentangling visual imagery and perception of real-world objects. Neuroimage 59:4064–4073.

Lorenz S, Weiner KS, Caspers J, Mohlberg H, Schleicher A, Bludau S, Eickhoff SB, Grill-Spector K, Zilles K, Amunts K (2015) Two New Cytoarchitectonic Areas on the Human Mid-Fusiform Gyrus. Cerebral Cortex 27:373–385.

Mahon BZ, Caramazza A (2011) What drives the organization of object knowledge in the brain? Trends in cognitive sciences 15:97–103.

Mahr JB (2020) The dimensions of episodic simulation. Cognition 196:104085.

Manning L (2000) Loss of visual imagery and defective recognition of parts of wholes in optic aphasia. Neurocase 6:111–128.

Mary A, Dayan J, Leone G, Postel C, Fraisse F, Malle C, Vallée T, Klein-Peschanski C, Viader F, Sayette Vdl, Peschanski D, Eustache F, Gagnepain P (2020) Resilience after trauma: The role of memory suppression. Science 367:eaay8477.

Marzi CA, Mancini F, Savazzi S (2009) Interhemispheric transfer of phosphenes generated by occipital versus parietal transcranial magnetic stimulation. Experimental brain research 192:431–441.

Mazard A, Laou L, Joliot M, Mellet E (2005) Neural impact of the semantic content of visual mental images and visual percepts. Brain Res Cogn Brain Res 24:423–435.

Mechelli A, Price CJ, Friston KJ, Ishai A (2004) Where bottom-up meets top-down: Neuronal interactions during perception and imagery. Cerebral Cortex 14:1256–1265.

Montagna S, Wager T, Barrett LF, Johnson TD, Nichols TE (2018) Spatial Bayesian latent factor regression modeling of coordinate-based meta-analysis data. Biometrics 74:342–353.

Moro V, Berlucchi G, Lerch J, Tomaiuolo F, Aglioti SM (2008) Selective deficit of mental visual imagery with intact primary visual cortex and visual perception. cortex 44:109–118.

Moulton ST, Kosslyn SM (2009) Imagining predictions: mental imagery as mental emulation. Philosophical transactions of the Royal Society of London Series B, Biological sciences 364:1273–1280.

O’Craven KM, Kanwisher N (2000) Mental imagery of faces and places activates corresponding stimulus-specific brain regions. Journal of cognitive neuroscience 12:1013–1023.

Owen AM, Coleman MR, Boly M, Davis MH, Laureys S, Pickard JD (2006) Detecting Awareness in the Vegetative State. Science 313:1402–1402.

Parkin BL, Ekhtiari H, Walsh VF (2015) Non-invasive human brain stimulation in cognitive neuroscience: a primer. Neuron 87:932–945.

Pearson DG, Deeprose C, Wallace-Hadrill SM, Burnett Heyes S, Holmes EA (2013) Assessing mental imagery in clinical psychology: a review of imagery measures and a guiding framework. Clinical psychology review 33:1–23.

Pearson J (2019) The human imagination: the cognitive neuroscience of visual mental imagery. Nature Reviews Neuroscience 20:624–634.

Pearson J (2020) Reply to: Assessing the causal role of early visual areas in visual mental imagery. Nature Reviews Neuroscience 21:517–518.

Pearson J, Naselaris T, Holmes EA, Kosslyn SM (2015) Mental Imagery: Functional Mechanisms and Clinical Applications. Trends in cognitive sciences 19:590–602.

Petersen SE, Posner MI (2012) The attention system of the human brain: 20 years after. Annual review of neuroscience 35:73–89.

Policardi E, Perani D, Zago S, Grassi F, Fazio F, Ladavas E (1996) Failure to evoke visual images in a case of long-lasting cortical blindness. Neurocase 2:381–394.

Riddoch MJ (1990) Loss of visual imagery: A generation deficit. Cognitive Neuropsychology 7:249–273.

Rode G, Cotton F, Revol P, Jacquin-Courtois S, Rossetti Y, Bartolomeo P (2010) Representation and disconnection in imaginal neglect. Neuropsychologia 48:2903–2911.

Rossi AF, Pessoa L, Desimone R, Ungerleider LG (2009) The prefrontal cortex and the executive control of attention. Experimental brain research 192:489–497.

Rossion B, Jacques C, Jonas J (2018) Mapping face categorization in the human ventral occipitotemporal cortex with direct neural intracranial recordings. Annals of the New York Academy of Sciences.

Rouder JN, Speckman PL, Sun D, Morey RD, Iverson G (2009) Bayesian t tests for accepting and rejecting the null hypothesis. Psychonomic bulletin & review 16:225–237.

Sack AT, Sperling JM, Prvulovic D, Formisano E, Goebel R, Di Salle F, Dierks T, Linden DE (2002) Tracking the mind’s image in the brain II: transcranial magnetic stimulation reveals parietal asymmetry in visuospatial imagery. Neuron 35:195–204.

Sadaghiani S, D’Esposito M (2015) Functional characterization of the cingulo-opercular network in the maintenance of tonic alertness. Cerebral Cortex 25:2763–2773.

Sasaoka T, Mizuhara H, Inui T (2014) Dynamic parieto-premotor network for mental image transformation revealed by simultaneous EEG and fMRI measurement. Journal of cognitive neuroscience 26:232–246.

Schacter DL, Addis DR, Buckner RL (2007) Remembering the past to imagine the future: the prospective brain. Nature Reviews Neuroscience 8:657–661.

Senden M, Emmerling TC, Van Hoof R, Frost MA, Goebel R (2019) Reconstructing imagined letters from early visual cortex reveals tight topographic correspondence between visual mental imagery and perception. Brain Structure and Function 224:1167–1183.

Seurinck R, de Lange FP, Achten E, Vingerhoets G (2011) Mental rotation meets the motion aftereffect: the role of hV5/MT+ in visual mental imagery. Journal of cognitive neuroscience 23:1395–1404.

Sheffield JM, Repovs G, Harms MP, Carter CS, Gold JM, MacDonald III AW, Ragland JD, Silverstein SM, Godwin D, Barch DM (2015) Fronto-parietal and cingulo-opercular network integrity and cognition in health and schizophrenia. Neuropsychologia 73:82–93.

Shelton AL, Pippitt HA (2006) Motion in the mind’s eye: comparing mental and visual rotation. Cognitive, Affective, & Behavioral Neuroscience 6:323–332.

Sirigu A, Duhamel J (2001) Motor and visual imagery as two complementary but neurally dissociable mental processes. Journal of Cognitive Neuroscience 13:910–919.

Soddu A, Boly M, Nir Y, Noirhomme Q, Vanhaudenhuyse A, Demertzi A, Arzi A, Ovadia S, Stanziano M, Papa M (2009) Reaching across the abyss: recent advances in functional magnetic resonance imaging and their potential relevance to disorders of consciousness. Progress in brain research 177:261–274.

Spagna A, Kim TH, Wu T, Fan J (2018) Right hemisphere superiority for executive control of attention. Cortex.

Spagna A, Martella D, Fuentes LJ, Marotta A, Casagrande M (2016) Hemispheric modulations of the attentional networks. Brain and Cognition 108:73.

Sunday MA, McGugin RW, Tamber-Rosenau BJ, Gauthier I (2018) Visual imagery of faces and cars in face-selective visual areas. PloS one 13:e0205041.

Thomas N, J., T. (1999) Are theories of imagery theories of imagination?: An active perception approach to conscious mental content. Cogntive Science 23:207–245.

Thorudottir S, Sigurdardottir HM, Rice GE, Kerry SJ, Robotham RJ, Leff AP, Starrfelt R (2020) The Architect Who Lost the Ability to Imagine: The Cerebral Basis of Visual Imagery. Brain Sci 10:59.

VanRullen R, Reddy L (2019) Reconstructing faces from fMRI patterns using deep generative neural networks. Communications Biology 2:193.

Wandell BA (2011) The neurobiological basis of seeing words. Annals of the New York Academy of Sciences 1224:63.

Wandell BA, Winawer J (2011) Imaging retinotopic maps in the human brain. Vision research 51:718–737.

Wang L, Mruczek RE, Arcaro MJ, Kastner S (2015) Probabilistic maps of visual topography in human cortex. Cerebral cortex 25:3911–3931.

Whittingstall K, Bernier M, Houde JC, Fortin D, Descoteaux M (2014) Structural network underlying visuospatial imagery in humans. Cortex 56:85–98.

Winlove CIP, Milton F, Ranson J, Fulford J, MacKisack M, Macpherson F, Zeman A (2018) The neural correlates of visual imagery: A co-ordinate-based meta-analysis. Cortex 105:4–25.

Wu T, Wang X, Wu Q, Spagna A, Yang J, Yuan C, Wu Y, Gao Z, Hof PR, Fan J (2019) Anterior insular cortex is a bottleneck of cognitive control. NeuroImage 195:490–504.

Xuan B, Mackie M-A, Spagna A, Wu T, Tian Y, Hof PR, Fan J (2016) The activation of interactive attentional networks. NeuroImage 129:308–319.

Yarkoni T, Poldrack RA, Nichols TE, Van Essen DC, Wager TD (2011) Large-scale automated synthesis of human functional neuroimaging data. Nature methods 8:665.

Yomogida Y, Sugiura M, Watanabe J, Akitsuki Y, Sassa Y, Sato T, Matsue Y, Kawashima R (2004) Mental visual synthesis is originated in the fronto-temporal network of the left hemisphere. Cereb Cortex 14:1376–1383.

Zago S, Corti S, Bersano A, Baron P, Conti G, Ballabio E, Lanfranconi S, Cinnante C, Costa A, Cappellari A (2010) A cortically blind patient with preserved visual imagery. Cognitive and behavioral neurology 23:44–48.

Zeman A, Dewar M, Della Sala S (2015) Lives without imagery – Congenital aphantasia. Cortex 73:378–380.

